# Tissue-wide cues are sensed at the cellular level to coordinate microtubule orientations in plants

**DOI:** 10.1101/2025.10.18.683221

**Authors:** Jordi Chan, Richard Kennaway, Jacob Newman, Enrico Coen

## Abstract

Plant morphogenesis depends on tissue-wide coordination of cortical microtubule orientations. This coordination is thought to depend on microtubules responding to mechanical stress orientations. Here we test this stress-sensing hypothesis by quantifying microtubule behaviours on different faces and edges of growing leaf cells. We show that microtubules orientations exhibit both cell-geometric and tissue biases. Stress sensing can account for tissue biases but not microtubule trajectories and densities. An alternative hypothesis is suggested by two edge behaviours: generation of microtubules within cell edges and a filter that prevents microtubules entering side faces at shallow angles. Incorporating these features into a combinatorial model in which cell face identities modulate microtubule behaviours, accounts for tissue coordination and observed dynamics. Thus, instead of orientations being sensed at the microtubule level, they are sensed at the cellular level through changes in statistical behaviours at cell faces and edges, coordinated across a tissue through cell polarity.

## Introduction

A fundamental problem in cell and developmental biology is how orientations of the cytoskeleton are coordinated between cells of a tissue. Such coordination plays a critical role in plants as the orientation of cortical microtubules guide cellulose synthesis and thus the extent to which cell walls yield to turgor and grow in different directions^1,2^. Cortical microtubules exhibit three features: *alignment*: microtubules tend to orient parallel with each other^3,4^, *orientation*: microtubule alignments preferentially orient parallel or perpendicular to cell-geometric or tissue axes^5–8^, and *persistent reorientation*: orientations may continually change over time^9–12^. Both orientations and the extent of persistent reorientation can vary between faces of the same cell^13–16^.

Alignment formation has been explained through microtubule-microtubule interactions. When the growing plus end of a microtubule encounters an obstructing microtubule, the plus end may undergo rapid depolymerisation (collision-induced catastrophe) with a probability that depends on angle of encounter^17^. In a population of microtubules, a chance excess of microtubules in one orientation leads to a higher survival probability for microtubules growing in that orientation, as they are less likely to undergo catastrophic collisions. This process reinforces the number of microtubules in that orientation, leading to alignment formation, through a process termed “survival of the aligned”^18–20^.

Orientation with respect to cell geometry has been explained through microtubule-edge interactions. If the probability of a microtubule undergoing catastrophe is enhanced at cell edges, either because of enhanced curvature or localised edge factors^21–24^, orientations are biased towards the long axis of the cell because microtubules travelling in that direction experience fewer edge encounters. Orientations biased towards the cell short axis have been explained by microtubules responding to mechanical stresses which can be higher in this direction^7,25–28^, or by microfibrils impeding microtubule growth^24^, or by microtubule-based nucleation ^29^.

Tissue-wide coordination of orientations has been explained through microtubules responding to mechanical stresses. The epidermis typically constrains growth of internal tissue, putting epidermal cell walls under tissue tension^30–32^. Cortical microtubules may be able to rotate or change their growth trajectories or growth rates to align with such tissue stresses^7,15,26,28,33^, leading to tissue-wide coordination.

Although many of the above hypotheses have been modelled^15,33–35^ the extent to which they can account for different microtubule behaviours and dynamics on faces of the same cell remains to be explored. A good test case is the leaf epidermal cell where the outer face exhibits little orientation preference while side faces show orientations perpendicular to the epidermal plane^15^.

Here we use photobleaching to track microtubule behaviours on different faces and edges of growing epidermal leaf cells. We show that microtubules on the outer face undergo persistent reorientation, with a cell-geometric bias towards the long axis of the cell. Both the outer and side faces also exhibit biases towards tissue axes: mediolateral for the outer face and orthoplanar for the side faces. To account for these observations, we develop a ground-state computational model based on quantified local microtubule interactions. Incorporating edge-induced catastrophe accounts for cell-geometric bias towards the long axis. Incorporating stress sensing accounts for tissue biases but predicted microtubule trajectories and side face densities are inconsistent with experimental data. An alternative hypothesis is suggested by behaviours observed at cell edges: generation of microtubules within the edge domain and a filter that prevents microtubules entering side faces at shallow angles. Incorporating these features into a combinatorial model in which different cell face identities modulate microtubule behaviours, accounts for both tissue-wide coordination and observed dynamics, including persistent reorientation. Our findings suggest that rather than orientational cues being sensed at the microtubule level, they are sensed at the cell level through cell face identities modifying statistical behaviours at cell edges. Cell face identities may be coordinated through tissue cell polarity systems, providing a general mechanism for tissue-wide coordination of microtubule and thus growth orientations in plants.

## Results

### Time-lapse imaging of growing leaf cells

To measure the orientation of microtubules, we imaged the abaxial surface (outer face) of growing leaf epidermal cells expressing TUA6-GFP (Tubulin tagged with GFP) at 20-30 min intervals (Figure 1A). For simplicity, we focused on the proximal region of the leaf primordium comprising cells with a near cuboidal shape (green dots Figure 1A). Cells outside this domain (pink dots) gave rise to meristemoid or pavement cell descendants at later stages of imaging.

**Figure 1.**
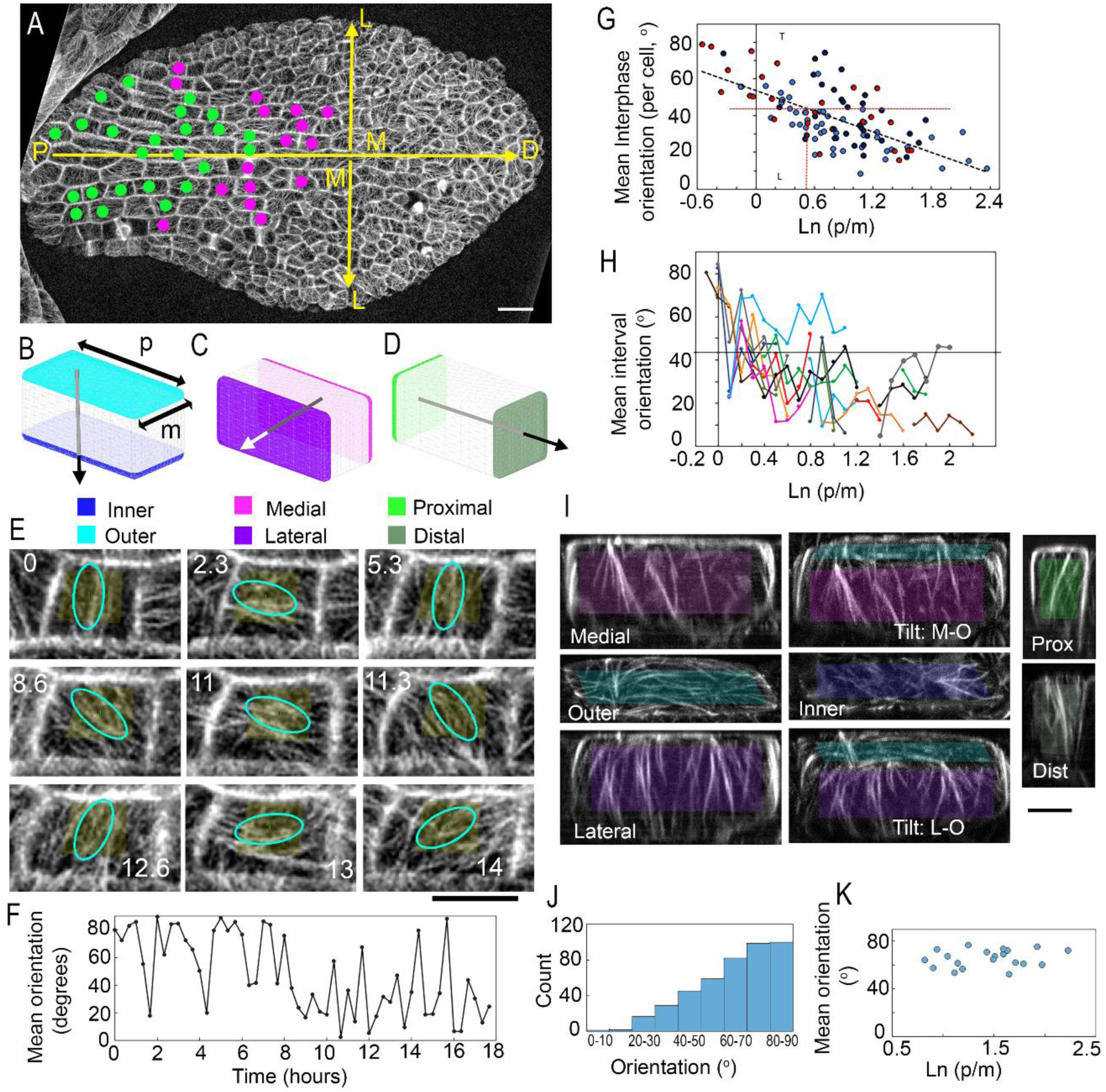
Arabidopsis leaf 1 at the start of imaging and labelling of cell faces. (**A**) Green dots indicate cells (and in some cases their daughters) included in the analysis. Pink dots mark cells that give rise to pavement cell and meristemoid descendants. Arrows indicate proximodistal and mediolateral axes. Primordium was approximately 165 µm wide at the start of imaging. Scale bar 20μm. (**B-D**) Faces and axes of leaf cells. (B) Outer (cyan) and inner (blue) faces perpendicular to the orthoplanar axis. Double-headed arrows = proximal length (p) and medial lengths (m) of cells (used to calculate p/m ratio). (C) Medial (magenta) and lateral (purple) faces perpendicular to the mediolateral axis. (D) Proximal (bright green) and distal (olive) faces perpendicular to the proximodistal axis. Arrows indicate polarity axes. (**E-F**) Dynamic alignments. (E) Movie frames showing microtubule orientation shifts during growth (time =hours). Ellipse indicate indicates mean orientation (width indicates coherency) of pixels in the region of interest (yellow box). (F) Mean orientation plotted against time for the cell shown above. (**G**) Mean interphase orientation plotted against mean ln (p/m) displays long axis geometric and ML biases. Data from 2 tracked leaves: light blue (50 cells; y = -16.58x + 47.903, R² = 0.52) and dark blue spots (32 cells; y = -20.978x + 60.318, R² = 0.30 ). Red dots indicate single-fluorescent cells (25 cells, 17 leaves, y = -21.141x + 57.587 ,R² = 0.60). Equation for all cells: y = -18.841x + 53.843, R² = 0.46. (**H**) Mean orientation on outer face of growing cells. Plot includes 20 individual cells changing Ln (p/m) ratios through growth. Many cells display negative slopes indicating alignments shift from transverse to longitudinal with increasing p/m ratios. The average number of movie frames per 0.1 interval was 12 (20 cells, 120 intervals, 1418 frames, stdev, 6.0). Average time per interval was 274 min (stdev = 132.5 min). (**I-K**) Side faces display orthoplanar bias. (I) Projections showing all 6 faces of a single-fluorescent cell. Transition between outer and side faces can be seen in tilted views: Tilt: M-O and Tilt:L-O. Bar = 5 µm. (**J**) Trajectories of microtubules on ML side faces (n = 20 cells, 434 trajectories). (**K)** Mean alignment is insensitive to Ln (p/m) ratio (n = 20 cells)

Cell edges were approximately aligned with three orthogonal tissue axes: (1) the proximodistal axis (PD axis), parallel to the midline of the leaf, (2) the mediolateral (ML) axis perpendicular to the midline, and the anticlinal or outer-inner (OI) axis, perpendicular to the plane of the leaf (Figure 1A). Cell faces perpendicular (normal) to these axes were named accordingly (Figure 1B-D). The cell division plane was typically oriented perpendicular to the PD axis. Cells grew about five times faster parallel to the PD axis (2.2% h^-1^, stdev = 0.64, 1 leaf, n=50 cells) than to the ML axis (0.46 % h^-1^, stdev = 0.41). Faster growth along the PD axis led to an increase in aspect ratio (PD length/ML length) until a cell division occurred, at which point aspect ratio was halved. Cells showed little growth parallel to the orthoplanar (OI) axis.

### Outer face microtubule orientations exhibit cell-geometric bias towards the long axis of the cell

Microtubule orientations on the outer abaxial face were highly dynamic, exhibiting persistent reorientation in many cases (Figure 1E-F). To determine whether the orientations were biased in relation to cell geometric or tissue axes, microtubule orientations on the outer face were quantified over entire interphases, excluding intervals around cell division (from formation of the preprophase band to ∼2h after division). Mean duration of interphases was 23h (stdev, 7.8 h, n=50) with measurements taken over 90% of the cell cycle duration at intervals of 20-30 mins. The prevalent microtubule orientation was measured at each time point relative to the PD axis (0° = parallel to PD axis). These values were then averaged over all time points for a cell, to give an interphase mean. At each time point the length along the PD axis (p, Figure 1B) and the length along the ML axis (m) were also measured, and the mean value of p/m calculated for the interphase.

Interphase means exhibited a negative correlation with ln (p/m) (Figure 1G, R^2^ = 0.36). Orientations were preferentially ML for short cells and PD for elongated cells Thus, microtubule orientation was biased towards the long axis of the cell. We also plotted mean orientations for individual cells over time. Increased ln (p/m) caused by growth correlated with decrease in angle, with a significant excess of negatively over positively sloped regression lines (Figure 1H).

Alignments on the inner face were sometimes visible in single fluorescent cells generated by photobleaching their neighbouring cells (Figure S1A-D). These examples showed (1) alignments on the inner and outer faces could differ in the same cell, (2) alignments on the inner face could differ between neighbouring cells and (3) alignments were dynamic (Figure S2). Thus, the inner face displayed persistent reorientation like the outer face but not in synchrony with it.

### Outer face microtubule orientations exhibit mediolateral tissue bias

If microtubule orientations on the outer face were purely dependent on cell geometry, a mean orientation of 45° would be expected for faces with a square shape (i.e. ln (p/m) = 0). However, the observed mean orientation for square faces was 54° ± 3.9 (intersection of regression line with vertical axis, Figure 1G). Mean orientation was 45° when ln (p/m) was 0.375 (vertical red dashed line), corresponding to p/m of about 1.45. Thus, in addition to a cell geometric bias, there was a tissue bias towards the mediolateral axis, with the two biases coming into balance when the cell was 45% longer than wide.

### Microtubule orientations on side faces exhibit orthoplanar tissue bias

To analyse microtubule orientations on side faces, we bleached cells neighbouring a cell of interest, allowing signal from its side face to be visualised (Figure S1A-D). To check whether bleaching affected microtubule behavior, we measured the orientation of microtubules on the outer face of the cell of interest and plotted it against ln (p/m) for 25 cells (Figure 1G and C, red dots). The points gave a similar distribution and regression equation to unbleached leaves, suggesting the bleaching the surround did not have a marked effect on microtubule behavior.

To see if microtubules displayed geometric/tissue bias on side faces, we measured their orientation of microtubules (Figure 1I). We focused on cells with proximodistal orientations on their outer faces. Microtubules on the medial and lateral side faces exhibited a mean orientation 65°, indicating an orthoplanar tissue bias (90° to the outer face) (Figure 1J). The orthoplanar bias was insensitive to p/m (Fig 1K). Microtubules on the proximal and distal side-faces also exhibited orthoplanar tissue bias (Fig 1I).

### A ground state computational model for microtubule dynamics

To understand the origins of geometric and tissue biases we first generated a ground state computational model of microtubule dynamics. To provide inputs for this model, we quantified local microtubule behaviours for the outer face using high time-resolution movies (5-8 sec) of cells marked with tubulin and EB1. Microtubule growth and depolymerization rates, were similar to those previously described (^36–38^, Table S1). Rescue of catastrophizing plus ends occurred spontaneously and often coincided with microtubule junctions (Figure S3A). We estimated a rescue probability for when a microtubule crossed another microtubule and in junction-free spaces (Table S1). New microtubules arose by branching either from the side of an extant microtubule (free branching, Figure S4A) or from the intersections of crossover junctions (Figure S4 B,C), as reported previously^39–42^. Free branching was biased towards the minus end (Table S1, Figure S4D,F), with 50 % (18/36) of branches initiated at or up to 0.5 µm away from the minus end. This minus-end bias was confirmed by treatments with low concentrations of oryzalin, which allowed individual microtubules to be tracked more easily (Figure S4E,G, Movie 1). In oryzalin-treated cells, 42 % (29/69) of microtubules were initiated within 0.5 µm of the minus end. The minus-end bias may explain fan-shaped clusters of branching microtubules previously reported for cells re-establishing their arrays following oryzalin treatment^43^. Formation of branched microtubule networks *in vitro* also arises from preferential nucleation near minus ends^44^.

When the growing plus end of a microtubule encountered an obstructing microtubule, the plus end had three possible fates: zippering, catastrophe or crossover. Low angle encounters favoured zippering, whereas higher angle encounters favoured crossover or catastrophe (Figure S3A,B). About 1 min after a crossover event, severing or branching could occur at the crossover junction (Figure S3C, Movie 2). Severing of either the crossing or obstructing microtubule was followed by catastrophe of the new plus end. New microtubules also arose by rescue of new plus ends (Figure S3C) – as previously reported^45^. Thus, some crossover-induced branching may correspond to severing followed by immediate rescue of the catastrophizing plus end.

The above behaviours were used as inputs into the ground state computational model. The surface of a cuboidal cell was initially populated with 100 randomly positioned and oriented microtubules, with plus ends growing along the cell surface and minus ends depolymerizing according to measured rates. Further nucleation was through free branching (from minus ends) and crossover-induced branching, each with probabilities and angle profiles based on observations (Figure S5D,F, Table S1). Rescue of depolymerizing ends was either spontaneous or crossover-induced, each with probabilities and angle distributions based on experimental measurements (Figure S5B, Table S1).

To capture the outcomes of microtubule-microtubule encounters, frequencies of events were binned into encounter angle intervals of < 20°, 20° - 40°, > 40° and converted into probabilities (Figure 2A). These probabilities were used to inform a decision tree for encounters (Figure 2B). Examples of different outcomes following encounters are illustrated in Movie 3 Each model was run for several hours, five times independently, for seven different p/m ratios.

**Figure 2.**
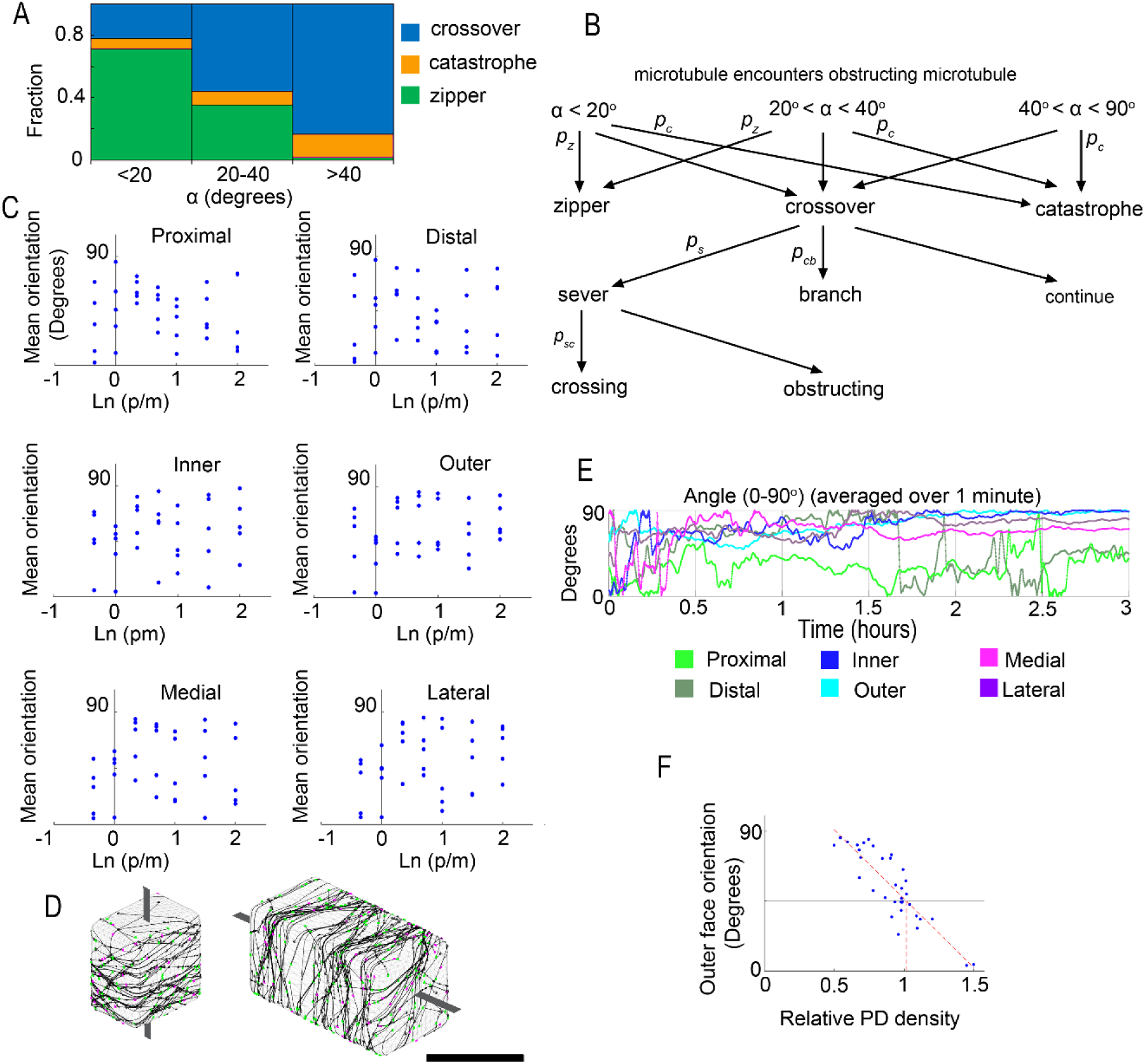
Microtubule behaviours in ground state model. (**A**) Histogram showing fractions of zipper, catastrophe or crossover upon encountering microtubules on outer face (n = 417, 9 cells). (**B**) Decision tree with probabilities for each type of outcome: *p_z_* = probability of zippering, *p_c_* = probability of catastrophe, *p_cb_* = probability of branching, *p_s_* = probability of severing, *p_sc_* = probability that it is the crossing microtubule that is severed. (**C**) Behaviour of alignments on faces of cells with different ln(p/m). Microtubules display random mean alignments on all faces irrespective of ln (p/m). (**D**) Snap shots of microtubule alignments at end of computer simulations in cells with p/m ratios of 0.7 and 2. Microtubules wrap around 4 faces along various axes. Dark grey lines indicate wrapping axes. (**E**) Dynamics of alignment formation on each face for an individual cell simulation. (**F**) Densities of microtubules on PD faces relative to the mean density on side faces for different simulations.

The ground state model led to randomly distributed mean orientations on all faces irrespective of p/m ratio (Figure 2C, D). For an individual cell, orientation fluctuated over time but typically settled after about an hour or so, with microtubules being co-aligned on four adjacent faces, wrapping around an axis (Fig 2D, E). The density of microtubules on PD faces, relative to the side face average, was higher when outer-face microtubules were aligned with the PD axis (0°) than with the ML axis (90°), as expected from the wrapping pattern (Figure 2F).

### Edge-induced catastrophe accounts for geometric bias towards the long axis of the cell

Previous computational modelling has shown that edge-induced catastrophe can lead to geometric bias towards the long axis of a cell^22,23^. To determine if microtubules exhibited edge-induced catastrophe, we made high resolution movies of single fluorescent cells with bleached neighbours. To track plus ends at edges we made ‘unfolded’ view movies by aligning 3 views: outer face (top-down view) – edge region (tilted view) – and side face (side view) (Figure S1E-H). To distinguish edge induced from microtubule-induced catastrophe, we focused on edges with relatively low microtubule densities, typically PD edges (Table S2). Microtubules often underwent catastrophe in the edge region (Figure 3A-C, Movie 4). We estimated an edge-induced catastrophe probability of about 0.7 for OI edges (25 microtubules, 17 catastrophes, 3 cells) and 1.0 for non-OI edges (43 microtubules, 43 catastrophes, 3 cells).

**Figure 3.**
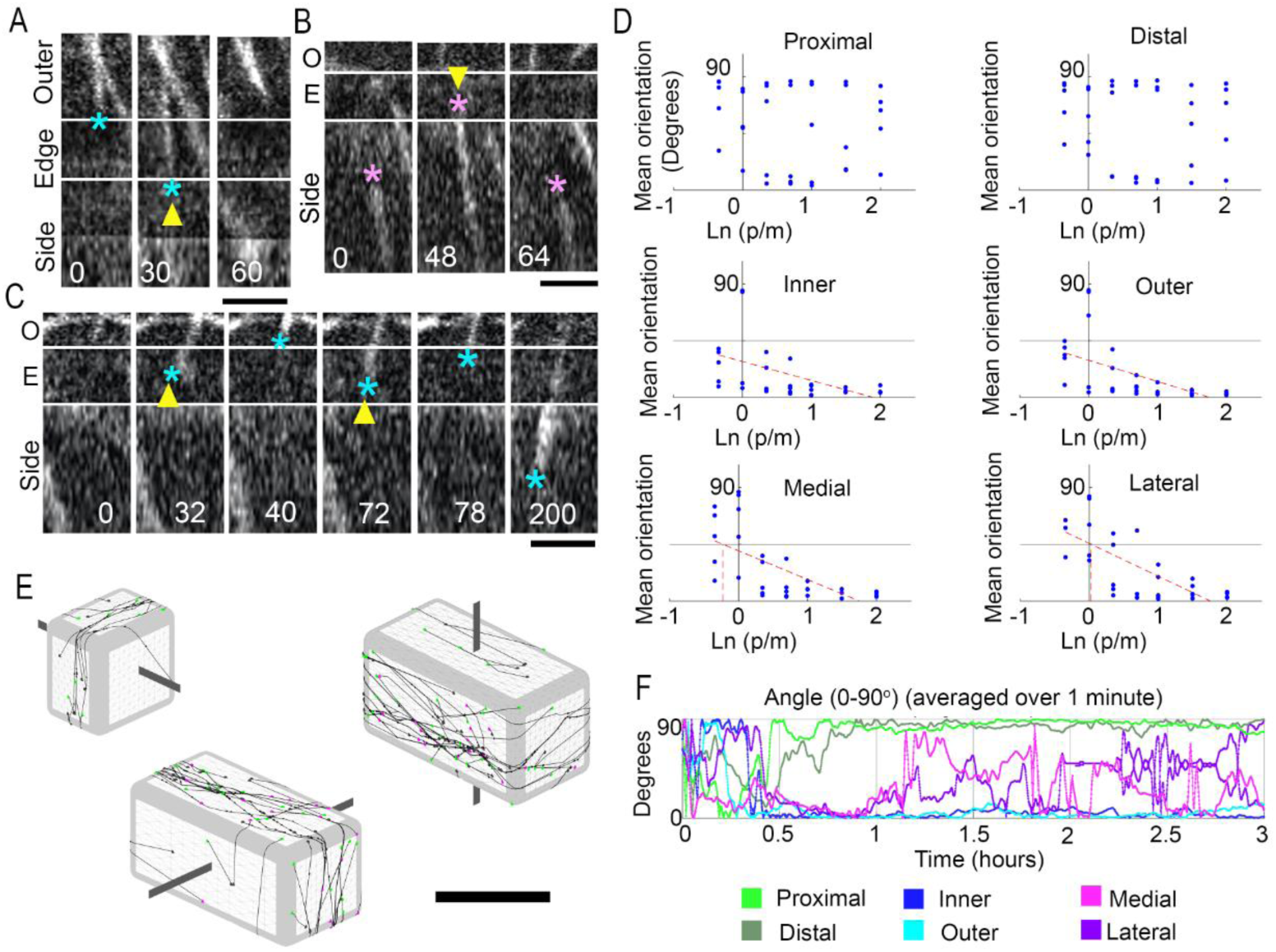
Edge-induced catastrophe models display geometric bias towards the long axis of the cell. (**A-B**) Frames from ‘unfolded’ view movie showing the behaviour of microtubules at edges. Microtubules displayed catastrophe when they arrived in the edge domain from either the outer face (A) or the side face (B). Coloured asterisks indicate plus end of microtubule at different times (time stamps shown in seconds). Yellow triangle indicates initiation of catastrophe. Bar = 2 µm. (**C**) Some microtubules underwent repeated catastrophe-rescue in the edge domain. (**D**) Behaviour of alignments on faces of cells with different ln(p/m) ratios. Alignments display long axis geometric bias on inner, outer, medial and lateral side faces. Line fit equations: Inner: y=-15x+28, R^2^=0.25; outer: y=-17x+29, R^2^=0.34, medial: -23x+40, R^2^=0.46; lateral=-26x+46, R^2^=0.54). (**E**) Microtubule alignments at end of runs in cells with p/m = 0.7 (left) or 2. Dark grey lines indicate wrapping axes. (**F**) Dynamics of alignment formation for a cell in which the wrapping axis is parallel to the mediolateral axis (p/m =2).

To determine whether this level of edge-induced catastrophe could account for the observed cell-geometric bias, we introduced it into the ground state model, assuming, for simplicity, that all edges had the same edge-induced catastrophe probability (0.7 for the minimum path length through the edge domain). Running this model gave cell-geometric bias in mean microtubule towards the long axis of the cell, evident from the negative slope of the mean angle vs ln (p/m) plot (Figure 3D). As with the ground-state model, microtubule orientation patterns tended to wrap around the cell. However, the wrapping axis was not randomly oriented. For short cells, the wrapping axis was parallel to the proximodistal axis, whereas for long cells the wrapping axis was parallel to either the mediolateral or orthoplanar axes (Figure 3E). Orientations became fixed on the wrapping faces but were more variable on the two remaining faces (Figure 3F). These results are consistent with the survival-of-the-aligned hypothesis as microtubules travelling along a cell’s long axis have higher survival probability.

### Stress-induced reorientation can account for tissue biases but not microtubule trajectories, densities or persistent reorientation

In the above model, ML faces of long cells often exhibited alignments parallel to the proximodistal axis (Figure 3 D-F), in contrast to the observed orthoplanar bias (Figure 1I-K). Orthoplanar tissue bias on side faces could be explained by microtubules responding to a tissue stress cue from the cell wall^15^. This cue could act by causing microtubules or their plus end trajectories to rotate to align with the direction of maximum stress^27^, or by accelerating plus end growth^33^. Microtubules on side faces showed no obvious rotation towards anticlinal (Movie 5), and plus end growth rates were similar to those on the outer face (Table S2). We therefore modelled the stress cue as causing the plus end trajectory to rotate towards orthoplanar as they travel through the side face. To account for mediolateral tissue bias on the outer face (Figure 1G), we also assumed a weaker mediolaterally-oriented stress cue on OI faces.

Introducing microtubule responses to these stress cues into the model with edge-induced catastrophe led to orthoplanar bias on side faces, and mediolateral bias on OI faces, matching the experimental observations (compare Figure 4A with Figure 1G and K). For shorter cells (ln (p/m) < 0.5), the wrapping axis was parallel to the proximodistal axis; whereas for longer cells the wrapping axis was parallel to the mediolateral axis (Figure 4B). Thus, observed tissue biases could be explained by a stress-cue model.

**Figure 4.**
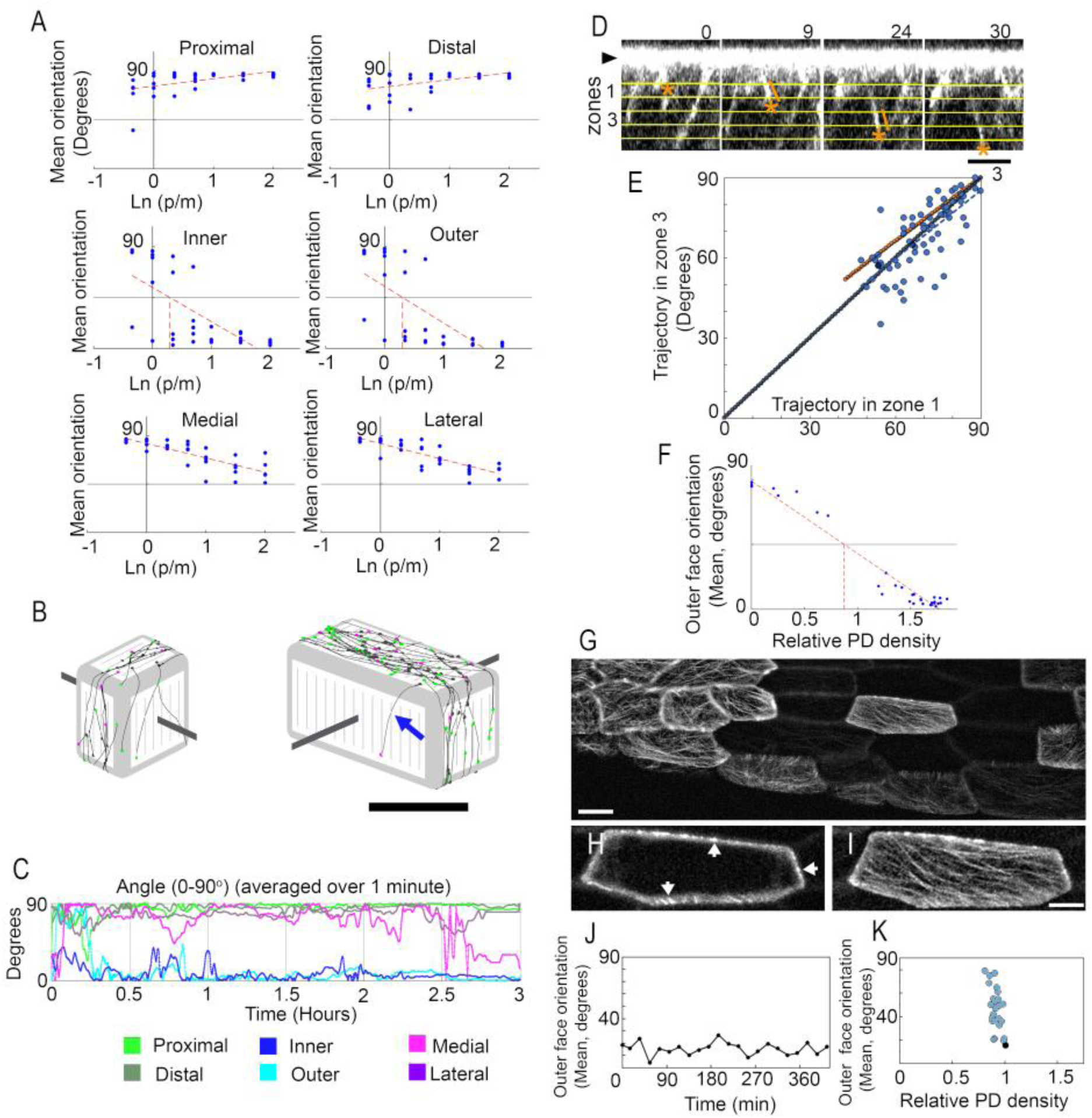
Stress-sensing model captures tissue biases but not microtubule trajectories or densities on side faces. (**A**) Mean microtubule orientations for different faces generated by the stress-sensing model, plotted against ln(p/m). Alignments display long axis geometric bias on inner and outer faces (strong negative slope of trend line), orthoplanar tissue bias on side faces (trend line above 45°), and mediolateral tissue bias on inner and outer faces (vertical red dotted line intersects ln(p/m) at value greater than 0). Line fit equations: Proximal: y=6.8x+76, R2=0.27; Distal: y=6.4x+76, R2=0.26; Inner: y=-30x+54, R2=0.49; Outer: y=-32x+55, R2=0.52, Medial: y=-13x+83, R2=0.65; Lateral: y=-14x+83, R2=0.71). (**B**) Microtubule patterns at end of runs in cells with p/m = 0.7 or 2. Thin grey lines indicate orientation of the stress field. Orientations wrap around four faces. Dark grey lines indicate wrapping axes. Blue arrow indicates a curved trajectory turning towards the stress cue. Bar = 10 µm. (**C**) Dynamics of mean microtubule orientations for a single cell with p/m=2. Orientations on outer and inner faces become parallel to the proximodistal axis, while those on the proximal and distal faces align with the orthoplanar axis. (**D**) Movie frames showing a microtubule entering side face. Trajectories were measured at entry, (zone 1, 0-1 µm deep) and later (zone 3, 2-3 µm deep). Orange asterisk indicates plus end. Yellow lines indicate zones. Bar = 3 µm. (**E**) Entry angle in zone 1 plotted against later angle in zone 3 for ML side faces (3 cells, 70 trajectories, 3 plants). Trajectories measured relative to horizontal edge. Lines indicate result if microtubules maintained the same trajectory (black ) or for turned towards the stress cue assuming same parameters as used in the above simulations (orange line, ). Number of points falling above (31) or below (35) the black trendline was not significantly different (Chi square, 1 degree freedom = 0.242, p = 0.6). The number of points falling above (17) or below (52) the orange trendline was significantly different (Chi square, 1 degree freedom = 17.8, p = 2.5 x 10^-5^). The experimental data gave rise to a trendline with the equation: y = 0.8554x + 8.5038 (R² = 0.5287). (**F**) Orientation of microtubules on outer face plotted against relative PD densities (microtubule density on PD faces divide by mean side face density), predicted by the stress-sensing model. Equation for line fit: y=-50x+89, R2=0.97. (**G-K**) Measuring relative side face density in single fluorescent cells. Example of cell surrounded by bleached neighbours (G). Bar = 10 µm. Same cell viewed as Z-projections of midplane (H) and the outer face (I). Arrows in H indicate cross sections of microtubules. Bar = 5 µm. (**J**) Projections of the outer face were used to plot mean orientation over time for a single cell. Longitudinal orientation is maintained in this example. (**K**) Densities of microtubules on PD faces relative to the mean density on side faces for single fluorescent cells. Pixel intensities were measured along PD and ML edges of midplane projections as a proxy for side face densities (n=25 cells, 17 leaves). Black spot indicates the cell shown in panels G-I. The clustered distribution differs from that predicted by the stress-sensing model (panel F).

The stress-cue model predicted curved trajectories for microtubules entering the side face obliquely (Fig 4B, arrowed), because of the time it took to align with the stress cue. To test whether trajectories curved, we tracked microtubules entering the ML side faces from the edge region, for cells showing PD orientations on the outer face (Movie 5). We plotted the angle at which plus end trajectories entered the side face (entry angle, zone 1, Figure 4D) against the trajectory angle further down the side face (later angle, zone 3) (microtubules that interacted with obstructing microtubules were ignored). The stress-cue model predicted that curved trajectory would give a difference between entry and later angles values, particularly for oblique entries (orange line, Figure 4E). The results were significantly different from this expectation (p = 2.5 x 10^-5^). The results were consistent with no significant difference between entry and later angles (black line, Figure 4E, p = 0.6).

Another prediction of the stress-cue model is two main wrapping patterns. Density of microtubules is relatively low when the wrapping axis is parallel to the proximodistal axis (Figure 4B, left), and high when the axis is perpendicular to this. This behaviour leads to a negative correlation between mean microtubule orientation on the outer face and mean microtubule density on PD faces relative to the side face mean (Figure 4F). To test this prediction, we measured TUA6-GFP intensities in the middle z-planes of side faces over extended periods of interphase (Figure 4 G,H). Densities on different side faces were integrated over time to obtain an average density for each side face. We also measured the mean orientation of microtubules on the outer face for each cell (Figure 4I-J). Mean microtubule density on PD faces relative to the side face mean was 0.8-1, irrespective of the orientation of the outer face (Figure 4K) which contrasts with the stress-cue model prediction (Figure 4F).

### Edge domains exhibit nucleation and edge-orthogonal filtering

In measuring entry angles into the ML side faces, we noticed a deficit of low angles despite the proximodistal orientation on the outer face (Figure 4F, note lack of datapoints for angles less than 50°). To explore the origin of this deficit, we imaged microtubules at edges of single fluorescent cells with proximodistal alignments on the outer face. Tracking microtubules arriving at edge domain from outer face showed that very few crossed on to the side face. Of 137 microtubules arriving at the edge domain (Fig 5B, cyan trajectories), only 4 (3%) travelled through the edge domain to reach the side face.

**Figure 5.**
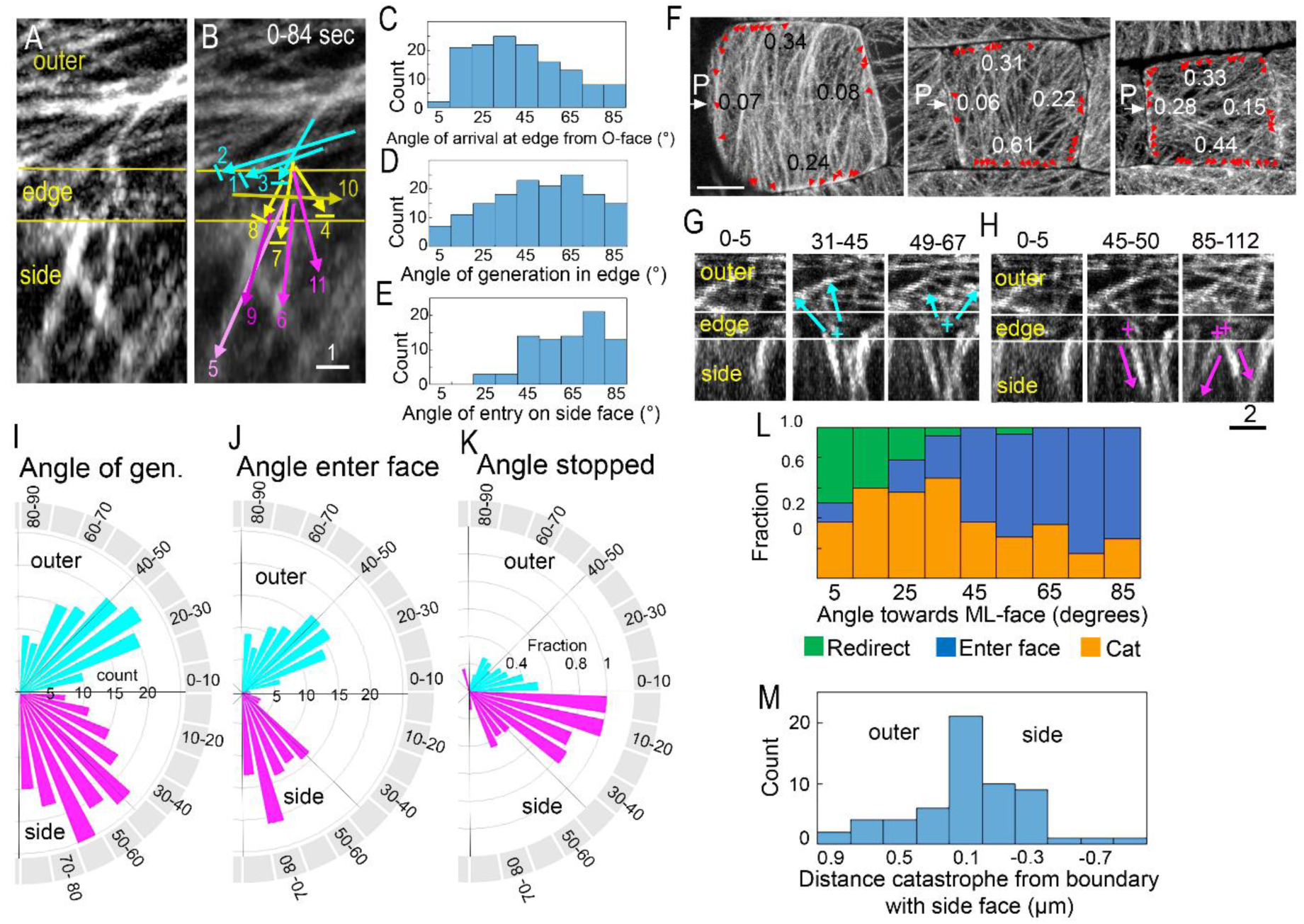
Generation and filtering of microtubules at cell edges. (**A-B**) Tracking microtubule trajectories across an edge. Time projection (84 sec) showing trajectories of microtubules on the outer face (outer), edge domain (edge) and side face (side). Some trajectories from the other face stop at the edge domain (cyan arrows). Microtubules generated within edge domain may stop before entering the side face (yellow arrows) or may enter the side face (magenta arrows). Microtubules were labelled in order of appearance. Short lines indicate catastrophe. Arrows indicate direction of plus end growth (See Movie 6 ).Yellow lines mark bounds of the edge domain (transition between outer and side face). Bar =1 µm. (**C**) Frequency histogram of trajectory angles arriving at edge domain from the outer face (e.g. cyan arrows in B, 2 cells,137 trajectories). Midpoint of alternate 10-degree-wide bins are numbered. (**D**) Frequency histogram of trajectory angles for microtubules generated within edge domain (e.g. yellow and magenta arrows in B, 2 cells, 153 trajectories). (**E**) Frequency histogram of trajectory angles for microtubules entering the side face (e.g. magenta arrows in B, 2 cells, 81 trajectories). (**F**) Emergence of microtubules from edges in square cells. Red arrowheads indicate sites of emergence from edges. P (white arrow) indicates the proximal face. Entry rates per µm of edge per min are shown by their corresponding edges. Bar = 4 µm. The emergence from ML and PD edges was significanly different from uniform (chi square value = 34, p=2.4 x 10^-7^, 2 degrees of freedom). (**G-H**) Movie frames of edge-generated microtubules that enter outer face (F, cyan arrows) or side face (G, magenta arrows). Crosses indicate start of trajectories, bar = 2 µm (See Movie 7). (**I**) Rose plot showing angle frequencies for edge-generated microtubules. Microtubules travelling towards outer face are in cyan (122 trajectories) and towards side faces are in magenta (153 trajectories), (2 cells, 275 trajectories). The distribution of angles towards each face was not significantly different from the mean distribution heading towards both faces (Chi square value = 3.8, p=0.15, 2 degrees of freedom). (**J**) Rose plot showing angle frequencies for edge-generated microtubules entering the outer face (cyan, 92 trajectories, 2 cells) or side faces (magenta, 81 trajectories, 2 cells). The distribution of trajectory angles entering each face was signifcantly different to the mean distribution entering the two faces (Chi square value = 11, p=0.004, 2 degrees freedom). (**K**) Rose plot showing the fraction of edge-generated microtubules blocked from entering the outer face (cyan) or edge (magenta). Calculated by dividing the number of stopped trajectories by the total number of trajectories genertated per bin. (**L**) : Cummulative fractions of edge-generated microtubules undergoing redirection (green), catastrophe (orange) or entrance onto the side face (blue) as a function of angle (2 cells, 154 trajectories). (**M**) : Frequency histogram of distance of catastrophe from the boundary with the side face (i.e. the lower yellow line in panel A, 2 cells, 59 trajectories). Midpoint of bins of 0.2 µm width are shown.

We also observed new trajectories generated within the edge domain, often in clusters of two or more microtubules (Fig 5B: yellow and magenta trajectores). Some of these trajectories, travelling at relatively steep angles, entered the side face (Fig 5B: magenta trajectories, compare generation trajectories in D with entry trajectories in E). Edge generation from OI edges was the main source of microtubules feeding the side faces, as branching from microtubules on side faces was rarely observed (Table S2), as was edge generation from non-OI (i.e. vertical) edges (No microtubules were observed departing vertical edges depite microtubules arriving at them (i.e. 43 arrivals, 3 movies). Whether edge-generated microtubules were nucleated *de novo* within the OI edge domains via branching, or from fragments of pre-existing microtubules that had undergone catastrophe or rescue was unclear.

Imaging emergence of microtubules from edges onto outer faces with an approximately square shape (chosen to avoid cell-geometric bias), revealed that average emergence rates were significantly higher along ML than PD edges (Figure 5F, 0.36 verses 0.14 per µm of edge per min, 4 cells, p=2.4 x 10^-7^), indicating a possible bias between these edges in rates of edge generation

Edge-generated microtubules travelled within the edge domain towards both outer and side faces (Fig 5G,H), with a symmetrical frequency distribution (Fig 5I). No significant difference was observed in the distribution of angles for trajectories of edge-generated microtubules heading towards outer face compared to the side face (p=0.15). By contrast, a significant difference was observed for trajectory angles entering side face compared with the outer face (Figure 5J p=0.004)). This entry asymmetry was caused by a significant depletion of microtubules entering the side face at shallow angles (0-30°, Figure 5K).

To understand the origin of deficit we followed the fate of edge-generated microtubules heading towards the side face. Edge-generated microtubules underwent redirection (plus end turns and grows along the edge) or catastrophe as they travelled through the edge domain. Measuring trajectory angles revealed their fate was angle dependent. Steep trajectories from 40-90 degrees had relatively high probabilities of entering the side face, whereas shallow trajectories displayed high probabilities for redirection or catastrophe, preventing their entrance (Figure 5L). Catastrophes tended to occurred close to the boundary with the side face (Figure 5M). These findings indicate that there is a filter close to the boundary with the side face that only allows microtubules arriving steep angles to pass through. We refer to this filter as an edge-orthogonal filter.

To explore the effect of an edge-orthogonal filter on microtubule orientations, we incorporated it into the ground state model. The filter was located at the junction between the edge domain of the OI faces and the side faces (blue lines, Figure 6A). Only microtubules arriving at this filter with angles of 50°-90°, travelling in either direction, could pass through. Those arriving at lower angles underwent catastrophe. Running this model yielded patterns in which microtubules wrapped around the longer edges (Figure 6A), corresponding to cell-geometric bias towards the short axis of OI faces (positive slope of regression line, Figure 6B middle panels) and orthoplanar tissue bias on the side faces (Figure 6B top and bottom panels). The preference for wrapping around longer edges may be explained because it favours survival by lowering the likelihood of oblique entry and thus catastrophe. Once established, a wrapping pattern was fixed (Figure 6C).

**Figure 6.**
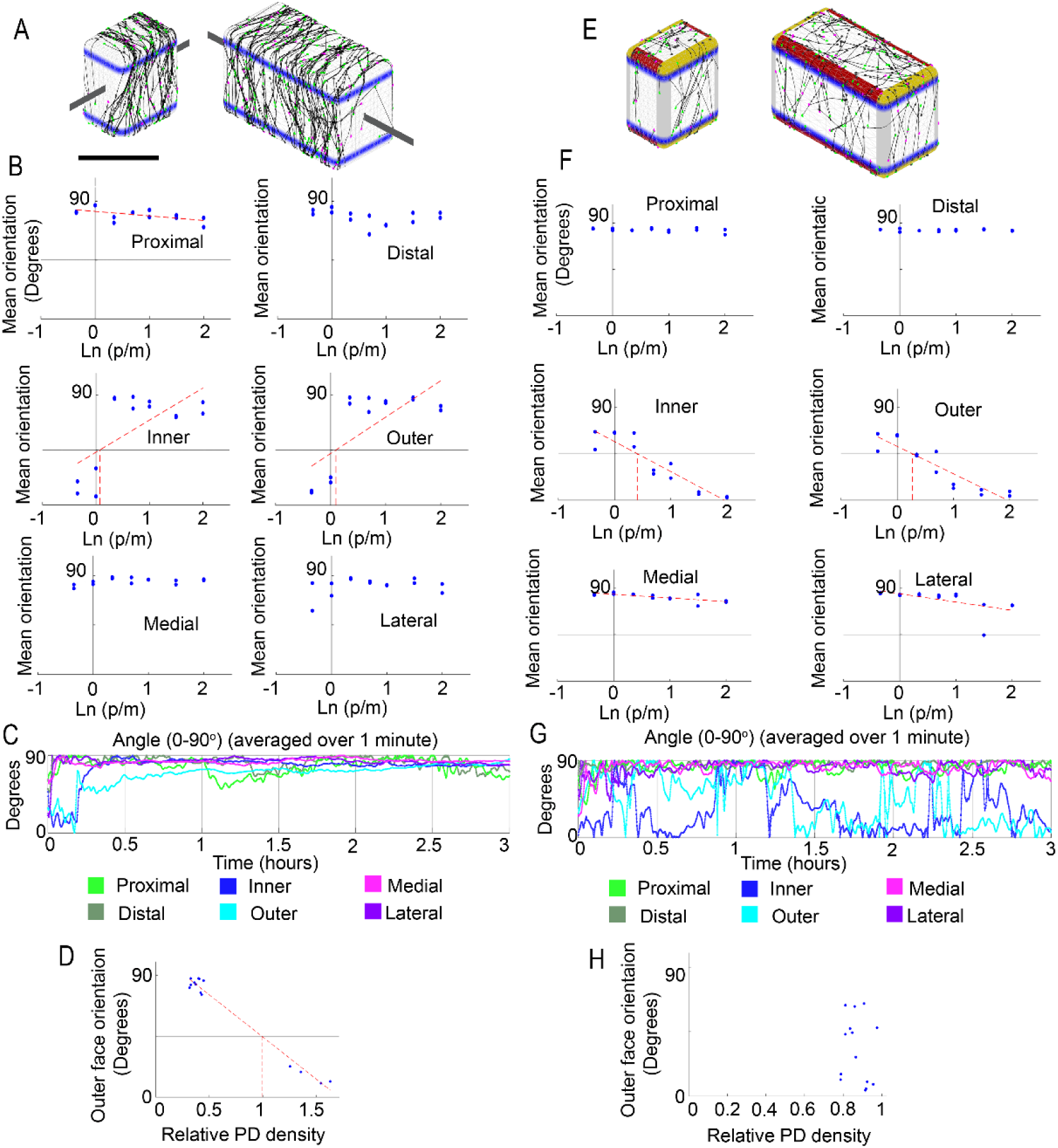
Outputs from models with edge-orthogonal filtering alone or in combination with other behaviours. (**A-D**) Results from incorporating edge-orthogonal filter alone into the ground state model. (**A**) Microtubule alignments at end of runs in cells with p/m = 0.7 or 2. Dark grey lines indicate wrapping axes. Bar = 10 µm. (**B**) Microtubule orientations on faces plotted against ln(p/m). Orientations on outer and inner faces display cell geometric bias towards the short axis of the cell. Line fit equations: Proximal: y=-3.5x+82, R^2^=0.35; Inner: y=26x+43, R^2^=0.45; Outer: y=35x+42, R^2^=0.54). (**C**) Dynamics of alignment formation. Alignments become fixed. Plot from the cell with Ln(p/m) of 2. (**D**) Relative PD densities of microtubules on side faces display cross feeding/wrapping. (Equation for line fit: y=-62x+1.1e+02, R2=0.98). (**E-H**) Combined model incorporating edge-induced catastrophe with face-modulated edge generation and filtering. (**E**) Snap shots of microtubule alignments at end of runs of combined model in cells with 0.7 and 2 Ln aspect ratio. Alignments do not wrap around 4 faces. (**F**) Simulations in cells with different ln(p/m) ratios. Alignments display long axis geometric bias and ML bias on outer and inner faces. All side faces display OI bias (Medial, Lateral, Proximal and Distal). 3 runs per cell aspect (Line fit: Inner: y=-29x+57, R2=0.8.5, Outer: y=27x+52, R2=0.84, Medial: y=-3.5x+84, R2=0.56. Lateral: y=-8x+85, R2=0.35). (**G**) Alignments display persistent reorientation on outer and inner faces. Plot from cell with Ln(p/m) of 2. (**H**) Relative PD densities of microtubules on side faces display clustered distribution.

Similar wrapping patterns were observed when edge generation (i.e. randomly oriented nucleation within the edge domain) was introduced into this model (Figure S6). Thus, with or without edge generation, an edge-orthogonal filter led to the opposite cell-geometric to edge-induced catastrophe.

In the above models we assumed that the edge-orthogonal filter only operates at OI edges. It is unclear whether it also operates at non-OI edges given the high catastrophe probability at these edges and their lack of edge-generation. An alternative model would be to have the orthogonal filter operating at all edges while also having a high catastrophe probability at non-OI edges, which would effectively neutralize the filter.

### Edge behaviours modulated by face identity lead to observed persistent reorientation, cell geometric and tissue biases

We next combined edge-generation and filtering with edge-induced catastrophe (Figure 4B). The combined model (Figure 6E-H) corresponds to the ground-state model with: (1) edge-induced catastrophe, (2) edge-orthogonal filtering, (3) OI edge generation, (4) higher edge generation at OI edges with ML than PD faces. Running this model gave orthoplanar tissue bias on the side faces (Figure 6G top and bottom) and cell-geometric bias towards the long axis of the OI faces (negative slope, Figure 6G middle).

Outer and inner faces also exhibited mediolateral tissue bias (vertical red dashed line intersects ln(p/m) above zero, Figure 6G middle), reflecting lower edge-generation rates from the OI-PD edges compared to the OI-ML edges. The model also gave persistent reorientation on the outer and inners faces, evident from the mean angle on these faces fluctuating over time (Fig 6F). The outer and inner face angles were not strongly correlated (compare dark blue and cyan lines), indicating that they reoriented independently of each other. Persistent orientation arose through the conflict between edge-induced catastrophe and edge-orthogonal filtering promoting opposite cell-geometric biases. At higher aspect ratios, the bias from edge-induced catastrophe was stronger, leading to overall geometric bias towards the long axis of the cell.

PD density was in the range 0.8-1.0, close to that observed experimentally (compare Figure 6H with Figure 4K). The reduced range in PD density compared that predicted by the stress-sensing model (Figure 4F) reflected OI-edge nucleation feeding microtubules to side faces rather than feeding coming solely from the OI faces.

The combined model therefore captured all key experimental observations: cell-geometric bias on the outer face towards the long axis of the cell, mediolateral tissue bias on the outer face, and orthoplanar tissue bias on the side faces, persistent reorientation on the outer face, no significant curving of microtubule trajectories on side faces, and PD density.

## Discussion

We show that microtubule orientations on different faces of growing leaf cells exhibit different behaviours. Those on the outer face exhibits persistent reorientation, with cell-geometric bias towards the long axis and mediolateral tissue bias. Those on the inner face reorient independently of the outer face. Those on side faces exhibit fixed orientations with orthoplanar tissue bias. The tissue biases can be explained through microtubule responding to stress cues, but this hypothesis is inconsistent with three observations: uncurved microtubule trajectories on side faces, proximal and distal microtubule densities relative to the mean side face density, persistent reorientation on the outer face.

To further understand the mechanisms underlying microtubule orientations, we imaged microtubule behaviours at edges of single fluorescent cells. In addition to previously described edge-induced catastrophe^21^, we observed generation of microtubules with the edge domain and an edge-orthogonal filter that blocks microtubules from entering the side face at shallow angles. We also observed enhanced emergence of microtubules from the medial and lateral edges of the outer face compared to the proximal and distal faces.

Incorporating these behaviours into a model showed that they could account for the observed tissue biases, persistent reorientation, uncurved trajectories and side face densities. The model involves modulation of microtubule behaviours through combinatorial interactions between three identities: edge (Figure 7A), OI face (Figure 7C) and PD face (Figure 7G) and. Incorporating edge-induced catastrophe alone into a ground-state model based on measured parameters leads to cell-geometric bias towards the long axis of the cell (Figure 7B), consistent with previous models^22,46^. Incorporating an edge-orthogonal filter at the OI identity border alone into the ground-state model generates cell-geometric bias towards the short axis (Figure 7D). This pattern is maintained by incorporation of OI edge generation (Figure 7E). Combining these behaviours with edge-induced catastrophe leads to persistent reorientation on the OI faces, together with cell-geometric bias towards the long axis (Figure 7F). Side face orientations are fixed and exhibit orthoplanar tissue bias. Introduction of reduced edge-generation at PD edges leads to the observed mediolateral tissue bias on OI faces (Figure 7H).

**Figure 7.**
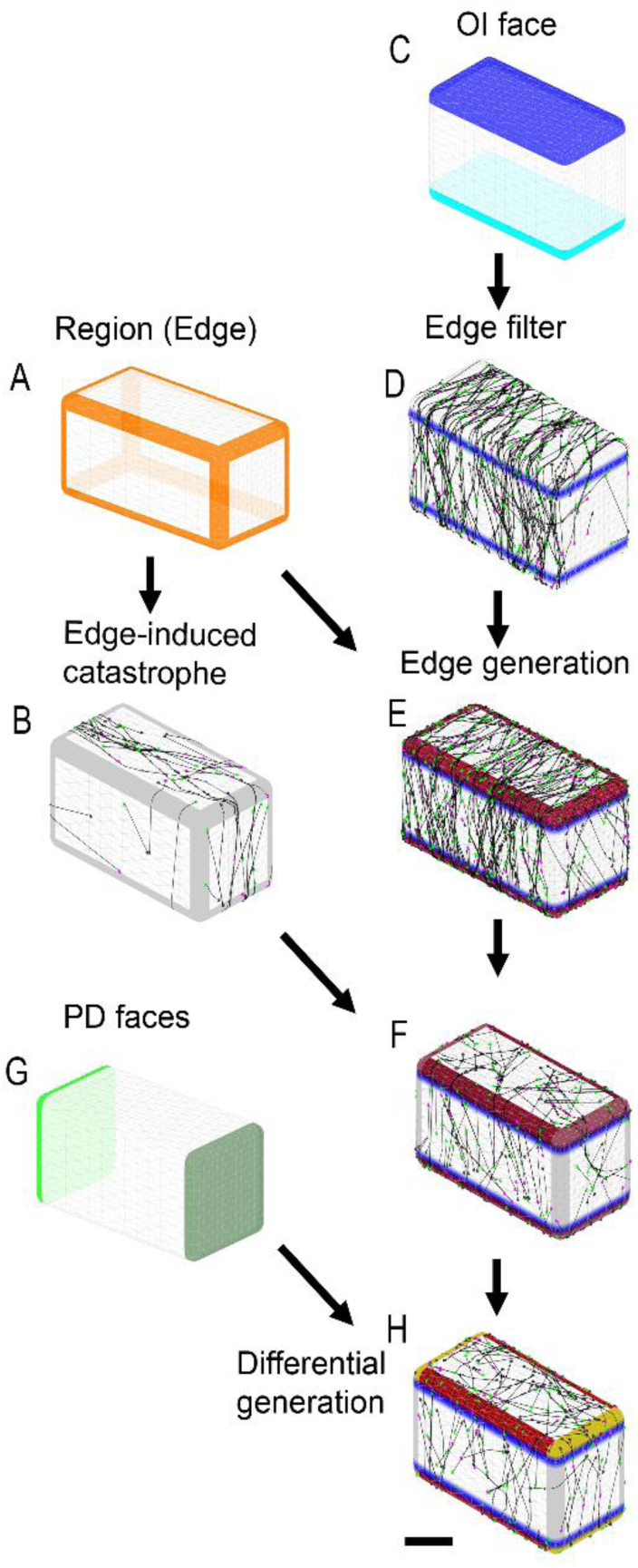
Combinatorial face and edge identity model to explain microtubule orientations. Edge region (**A**) can induce catastrophes leading to geometric bias towards the long axis of the cell (**B**). OI face identity (**C**) can specify location of an edge-orthogonal filer at its outer limits which leads to geometric bias towards the short axis of the cell (**D**). OI identity in combination with edge region identity can enhance generation from OI edges (**E**). Combining models B and E leads to persistent reorientation on OI faces with geometric bias towards the long axis, together with orthoplanar tissue bias on the side faces (**F**). PD face identity (**G**) can reduce edge generation to produce mediolateral tissue bias on the OI faces (**H**).

The combinatorial model depends on three identities: edge, PD face, and OI face. Edge effects may arise through higher curvature of edges^21^ and/or proteins that localize to edge domains^21,47^. Evidence for PD face identities comes from localization of proteins such as BASL and PIN^48,49^. In the case of BASL, interactions with microtubules have been proposed for stomatal lineage cells^50,51^. However, as the leaf cells analysed here do not form stomata, other polarity proteins would likely be involved. There is currently no molecular evidence for OI identity in leaf cells, although OI-polarised proteins have been observed for roots which have radial symmetry^52^. OI polarity for leaf cells seems likely given the greater thickness of outer epidermal cell walls compared to inner walls^53,54^.

By guiding cellulose synthesis, the microtubule patterns generated by the combinatorial model may account, at least in part, for the principal directions of epidermal cell growth. Orthoplanar microtubule orientation on side faces would promote microfibril alignments that reduce growth rate in epidermal cell thickness while allowing growth in side wall length, consistent with experimental observations. Persistent microtubule reorientation on the OI faces would produce no major direction of microfibril orientation, consistent with greater growth rate in the epidermal plane. Transverse tissue bias on the OI faces would promote greater planar growth rate parallel than perpendicular to the proximodistal axis, as observed. However, the transverse bias is weak, whereas planar growth is about five times faster parallel to the proximodistal axis than perpendicular to it. This discrepancy may be resolved if epidermal growth is partly driven by anisotropic growth of internal tissue which causes tissue stress parallel to the proximodistal axis^13,14,55^.

The combinatorial model proposed here contrasts with stress-sensing models. Instead of an orientational cue being sensed by microtubules, tissue biases arise through modulation of microtubule behaviours by cell face identities (OI and PD). Tissue biases are therefore a statistical outcome of polarities across a cell rather than a response of microtubules to a local orientational cue. Separating the role of mechanical stresses and tissue cell polarity in controlling microtubule orientation has been difficult because the two are often correlated with each other^32^. The most compelling data in support of the stress-sensing hypothesis comes from mechanical perturbations, such as squeezing of meristems which can cause weak but statistically significant alignments with the direction of maximal stress^26^. However, the observation of cell-geometric bias towards the long axis reported here and elsewhere^21–23^, may explain these results through changes in cell geometry (e.g. squeezing of cell shape) brought about by such mechanical perturbations.

Although the combinatorial face identity model proposed here is consistent with experimental observations, it begs many questions. First, it is unclear how edge generation occurs. It could be *de novo* or through branching and/or rescue of microtubule fragments generated by catastrophes in edge regions^56^. It is also unclear how the edge-orthogonal filter works. This filter operates in the opposite manner to curvature-induced catastrophe ^21^ as it blocks trajectories with low angles and thus low curvature. One possibility is that the filter provides a mechanical barrier such that only microtubules that encounter the barrier at steep angles have a sufficient force component to cross the barrier^57^. Other possibilities are that curved trajectories recruit growth survival factors^58^, or that oriented proteins localized at the border prevent transits at oblique angles.

A further question is how such a model might explain microtubule orientations in other tissues. For example, in hypocotyl epidermal cells the inner face exhibits fixed orientations orthogonal to the stem axis, while the outer face exhibits persistent reorientation^13,14^. This observation could be explained by inner face identity reducing edge-induced catastrophe. Interior cells of hypocotyls and roots exhibit fixed microtubule orientations orthogonal to the main growth (PD) axis^13,14,59,60^. This orientation might be explained by orthogonal filtering at cell edges parallel with the PD axis. Switches in orientation following environmental signals, such as light^61,62^ or hormones^63,64^, may involve changes in the relative contribution of different face or edge-modulated behaviours. High resolution imaging and quantification of microtubule dynamics at cell edges and interactions with polarity proteins may help resolve these questions.

## Materials and Methods

### Plant material and growth conditions

Seeds were sterilised with chlorine gas for 16 hours. The reaction was initiated by adding 5 ml of 37 % hydrochloric acid to 100 ml of a 10 % (w/v) sodium hydrochlorite solution.

Gas-sterilised *Arabidopsis thaliana* seeds were sown on plates containing 1 % (w/v) Agar, 1 % (w/v) glucose, 0.43 % (w/v) Murashige & Skoog powdered medium including vitamins, 3 mM MES, PH 5.7. Plates were stratified at 4 °C for 2 days and then grown at 20 °C under a 16 hour light/8 hour dark cycle.

6-day old seedlings were transferred from plates into a live-imaging chamber^68,69^. The chamber was perfused with 0.43 % (w/v) Murashige & Skoog powdered medium including vitamins containing 1 % (w/v) glucose, pH 5.8. Plants contained GFP-reporters for either tubulin and/or EB1^70–72^. High time resolution movies (∼3-8 sec intervals) of microtubule dynamics were taken of Arabidopsis Col-0 seedlings containing GFP-TUB6 and EB1-GFP. Extended imaging of microtubule arrays and side face densities were carried out using seedlings expressing TUA6-GFP.

For some experiments, the density of microtubule arrays was reduced by adding oryzalin^73^. 3.2 µM oryzalin was added to the chamber 3 days after plants had been transferred into the chamber. Leaves were imaged the next day.

### Confocal microscopy

Extended live imaging was carried out using a Zeiss Exciter confocal laser scanning microscope equipped with a x40/1.3 NA objective lens. Confocal sections were collected at a z-spacing of 0.7 µm and z-stacks acquired with an interval of 3-30 mins. Higher resolution movies of the outer face were imaged using a VisiTech (UK) spinning disc confocal microscope equipped with a 100 x/1.4 NA objective lens. Confocal sections were collected at a z-spacing of 0.3-0.5 µm and z-stacks acquired with an interval of 5-7.5 sec. Fluorescence was detected using a Hamamatsu Orca ER cooled CCD camera with 1x binning, set a 1.0 sec exposure time. GFP was excited using the 488 nm line of an argon laser and emitted light filtered through a 500-550 nm band-pass.

### Generating and imaging single fluorescent cells

Single-fluorescent cells were generated by bleaching fluorescence (TUA6-GFP or TUB-GFP - and EB1-GFP) in their neighbouring cells using a Zeiss (Exciter or 780 or 880) confocal laser scanning microscope equipped with a x40/1.3 NA, x63/1.4 NA or x100/1.5 NA objective lens. Fluorescence was specifically beached in neighbouring cells by targeting the laser on to their outer faces using the region of interest tool. To avoid bleaching the cell of interest, the regions avoided its edges (Figure S1B). When specifically bleached, fluorescence was only observed along edges of the cell-of interest and not between interfaces of bleached cells (Figure S1C,D). We assume that targeting the cortex results in fluorescence being removed on all cell faces because TUA6-GFP moves onto the outer face after being released by microtubule turnover.

Experiments were carried out on the first leaf of 7-10 day old seedlings. Older cells (as shown in Figure 3A-C) were obtained from 13-16 day old seedlings.

High resolution imaging of outer and side faces of single fluorescent cells was carried out using Zeis 780 and 880 confocal microscopes, fitted with x40/1.3 NA objective lenses. A box capturing the edge-of-interest and part of the outer face was selected using the region-of-interest tool. To be able to track microtubules, the size of the box was limited by an acquisition speed of 1.5-3 secs to collect a z-thickness of up to 5 µm, with a step size of 0.2 µm. Bidirectional scanning, no averaging and maximum scan speeds were used to help acquire z-stacks within the time limit.

### Image processing

All processing of z-stacks/images was carried out using Fiji, a version of ImageJ^74^. Tools included in this software were used to project z-data using maximum or sum projection, rotate, crop, align, reslice, adjust brightness and contrast, background subtract, and add scale bars and time stamps to movies. Montages were assembled using Adobe Photoshop. Some movies were processed to reduce noise, which involved sum projection using the Group Z-projector tool followed by Rolling-ball background subtraction. Movies were then aligned using the StackReg plugin^75^.

Mean microtubule alignments were measured from movies of the outer-epidermal face using the OrientationJ plugin of Fiji^76^. Angles of microtubules on side faces were measured manually because OrientationJ pluggin was sensitive to the z-step spacing used to collect the images.

All angles were measured using the angle tool of Fiji. Branching angles were measured relative to the growing plus end of the mother microtubule or relative to the crossing microtubule if birth occurred at a junction. The trajectory of rescue was measured relative to the original direction of growth.

### Mean interphase orientation

The cell of interest was cropped from the time-lapse data and then one of its edges (proximal or distal) aligned between movie frames. Movies were cropped to remove any biases arising from cell division – i.e. 20-30 mins before cortical microtubules started to bunch together into the pre-prophase band and 2 hours after division, when the cortical array establishes through microtubules growing from edges of the new cell wall.

A region of interest was then selected inside the cell using polygonal tool of ImageJ. The ROI was drawn parallel with PD axis of leaf and avoided cell edges. Mean microtubule orientation in each frame was then measured using OrientationJ (measure tool) pluggin of of Fiji. The mean orientation values were imported into excel and used to calculate mean interphase orientation. Print screen was used to copy images of cells with ellipses shown in Figure 1E.

### Cell shape

Cell outlines labelled by TUA6-GFP were used to quantify cell geometry. Polygonal cells tended to have 4 edges: 2 edges parallel with PD axis and 2 edges parallel with ML axis (*p* and *m*, Fig. 1B). The length of these edges was measured using the Line tool of Fiji and the average values of *p* and *m* calculated.

To assign a single value of *p*/*m* to growing cells, averages of *p* and *m* were taken from 2 measurements taken at the start and end of the movie. Values of *p* and *m* between these time intervals were interpolated to generate plots of Ln(*p/m*) versus mean orientation. Ln(*p/m*) was split into 0.1 bins and the mean orientation of microtubules estimated per bin.

### Growth rates

PD and ML growth rates were estimated by taking (ln final length - ln initial length) / time (min). Growth rates were expressed as % per hour (i.e. growth rates per min were multiplied by 60 to convert to µm per hour and then by 100 to give % per hour).

### Relative PD density

Pixel intensities were measured along edges of midplane projections from single-fluorescent cells as a proxy for side face densities. The mean pixel intensity was measured for proximal, distal, medial and lateral edges every 5-10 frames using line tool of imageJ. Average pixel intensities were then calculated for PD (sum of proximal and distal edges) and ML (sum of medial and lateral edges) sides of the cell. The relative PD density was calculated as 2x PD/ (ML+ PD).

### ‘Unfolded’ movies

‘Unfolded’ movies of single-fluorescent cells were made by combining planar views of the outer and side faces and a tilted view of the edge domain, using the combine tool of Fiji. The side face view and tilted views were projected using reslice and 3D project tools of Fiji, respectively.

All angles of new EB1 comets starting in the edge domain were measured in the Xtilt view. It was not possible to distinguish new comets generated by either branching, rescue, sever-rescue, break-rescue or turning/lateral movements. Thus, microtubules obviously entering the edge domain from the side face or outer face were ignored – since these trajectories did not start in the edge domain. Trajectories starting in edge domain that resulted from rescue of microtubules in the edge domain were included as it was too difficult to separate rescue from nucleation.

Angles of trajectories starting and ending in edge domain were measured in the Xtilt view. Some trajectories started in edge domain and ended on the side- or outer face, without deviation. These were measured as a single trajectory from their site of origin to a point on the adjacent face. Some trajectories changed angle on entering the side face or the outer face (due to turning through zippering or curvature). These cases were measured twice – first in the Xtilt view up until the boundary with the adjacent face and then from the boundary to a point on the side face or outer face. The Xtilt measurement was used to plot trajectories starting in edge domain. The side or outer face measurements were used to plot trajectories (entering) on the side or outer face, respectively.

Some trajectories started close to the boundary with an adjacent face – these were measured from the boundary to a point on the side face/outer face (i.e. these trajectories being too short to measure in the Xtilt view alone). These measurements were used to plot trajectories starting in the edge domain and plots of trajectories (entering) on the side or outer face respectively. In addition, distances of catastrophe/origin of new trajectories were made relative to the boundary of the edge domain with the side face.

## Acknowledgments

We thank Andrew Bangham (1947-2014) for ideas and inspiration. We thank Anna Hanner, Daniel Cosgrove, Pauline Durand, Sulin Zhang, Desmond bradley and members of the Coen lab for helpful discussions. We thank Grant Calder and the JIC BioImaging facility for helping develop confocal microscopy techniques. We thank JIC Horticultural services for growing and looking after plants.

## Notes

### Competing Interest Statement

The authors have declared no competing interest.

## References

1. Green, P.B. (1962). Mechanism for Plant Cellular Morphogenesis. Science (1979) 138, 1404–1405. 10.1126/science.138.3548.1404.

2. Paredez, A.R., Somerville, C., and Ehrhardt, D. (2006). Visualization of Cellulose Synthase with Microtubules. Science (1979) 312, 1491–1495. 10.1126/science.1126551.

3. Ledbetter, M.C. and Porter, K.R. (1963). A ‘microtubule’ in plant cell fine structure. J. Cell Biol. 19, 239–250.

4. Hardham, A.R., and Gunning, B.E.S. (1977). The length and disposition of cortical microtubules in plant cells fixed in glutaraldehyde—Osmium tetroxide. Planta 134, 201–202.

5. Durand-Smet, P., Spelman, T.A., Meyerowitz, E.M., and Jönsson, H. Cytoskeletal organization in isolated plant cells under geometry control. 10.17863/CAM.51754.

6. Chan, J. (2012). Microtubule and cellulose microfibril orientation during plant cell and organ growth. J. Microscopy 1, 23–32. 10.1111/j.1365-2818.2011.03585.x.

7. Colin, L., Chevallier, A., Tsugawa, S., Gacon, F., Godin, C., Viasnoff, V., Saunders, T.E., and Hamant, O. Cortical tension overrides geometrical cues to orient microtubules in confined protoplasts. 10.1073/pnas.2008895117/-/DCSupplemental.

8. Oda, Y. (2015). Cortical microtubule rearrangements and cell wall patterning. Front Plant Sci 6, 236. 10.3389/fpls.2015.00236.

9. Mayumi, K., and Shibaoka, H. (1996). No TitleThe cyclic reorientation of cortical microtubules on walls with a crossed polylamellate structure: effects of plant hormones and an inhibitor of protein kinases on the progression of the cycle. Protoplasma 195, 112–122.

10. Hejnowicz, Z. (2005). Autonomous changes in the orientation of cortical microtubules underlying the helicoidal cell wall of the sunflower hypocotyl epidermis: Spatial variation translated into temporal changes. Protoplasma 225, 243–256. 10.1007/s00709-005-0091-9.

11. Chan, J., Calder, G., Fox, S., and Lloyd, C.W. (2007). Cortical microtubule arrays undergo rotary movements in Arabidopsis hypocotyl epidermal cells. Nat Cell Biol 9, 171–175. 10.1038/ncb1533.

12. Thoms, D., Vineyard, L., Elliott, A., and Shaw, S.L. (2018). CLASP facilitates transitions between cortical microtubule array patterns. Plant Physiol 178, 1551–1567. 10.1104/pp.18.00961.

13. Chan, J., Eder, M., Crowell, E.F., Hampson, J., Calder, G., and Lloyd, C. (2011). Microtubules and CESA tracks at the inner epidermal wall align independently of those on the outer wall of light-grown Arabidopsis hypocotyls. J Cell Sci 124, 1088–1094. 10.1242/jcs.086702.

14. Crowell, E.F., Timpano, H., Desprez, T., Franssen-Verheijen, T., Emons, A.M., Höfte, H., and Vernhettes, S. (2011). Differential regulation of cellulose orientation at the inner and outer face of epidermal cells in the Arabidopsis hypocotyl. Plant Cell 23, 2592–2605. 10.1105/tpc.111.087338.

15. Zhao, F., Du, F., Oliveri, H., Zhou, L., Ali, O., Chen, W., Feng, S., Wang, Q., Lü, S., Long, M., et al. (2020). Microtubule-Mediated Wall Anisotropy Contributes to Leaf Blade Flattening. Current Biology 30, 3972–3985.e6. 10.1016/j.cub.2020.07.076.

16. Yuan, M., Warn, R.M., Shaw, P.J., and Lloyd, C.W. (1995). Dynamic microtubules under the radial and outer tangential walls of microinjected pea epidermal cells observed by computer reconstruction. The Plant Journal 7, 17–23. 10.1046/j.1365-313X.1995.07010017.x.

17. Dixit, R., and Cyr, R. (2004). Encounters between Dynamic Cortical Microtubules Promote Ordering of the Cortical Array through Angle-Dependent Modifications of Microtubule Behavior. Plant Cell 16, 3274–3284. 10.1105/tpc.104.026930.

18. Tindemans, S.H., Hawkins, R.J., and Mulder, B.M. (2010). Survival of the aligned: Ordering of the plant cortical microtubule array. Phys Rev Lett 104, 1–4. 10.1103/PhysRevLett.104.058103.

19. Baulin, V.A., Marques, C.M., and Thalmann, F. (2007). Collision induced spatial organization of microtubules. Biophys Chem 128, 231–244. 10.1016/j.bpc.2007.04.009.

20. Saltini, M., and Deinum, E.E. (2024). Microtubule simulations in plant biology: A field coming to maturity. Curr Opin Plant Biol 81, 102596. 10.1016/j.pbi.2024.102596.

21. Ambrose, C., Allard, J.F., Cytrynbaum, E.N., and Wasteneys, G.O. (2011). A CLASP-modulated cell edge barrier mechanism drives cell-wide cortical microtubule organization in Arabidopsis. Nat Commun 2, 430. 10.1038/ncomms1444.

22. Chakrabortty, B., Blilou, I., Scheres, B., and Mulder, B.M. (2018). A computational framework for cortical microtubule dynamics in realistically shaped plant cells. 1–26.

23. Mirabet, V., Krupinski, P., Hamant, O., Meyerowitz, E.M., Jönsson, H., and Boudaoud, A. (2018). The self-organization of plant microtubules inside the cell volume yields their cortical localization, stable alignment, and sensitivity to external cues. PLoS Comput Biol 14, 1–23. 10.1371/journal.pcbi.1006011.

24. Tian, T.Y.Y., Wasteneys, G.O., Macdonald, C.B., and Cytrynbaum, E.N. (2024). Conflicting roles of cell geometry, microtubule deflection and orientation-dependent dynamic instability in cortical array organization. Preprint, 10.1101/2024.09.07.611822 https://doi.org/10.1101/2024.09.07.611822.

25. Richmond P, A. (1983). Patterns of cellulose microfibril deposition and rearrangement in Nitella: In vivo analysis by a birefringence index. J. Appl. Polym. Sci. 37, 107–122.

26. Hamant, O., Heisler, M.G., Jönsson, H., Krupinski, P., Uyttewaal, M., Bokov, P., Corson, F., Sahlin, P., Boudaoud, A., Meyerowitz, E.M., et al. (2008). Developmental patterning by mechanical signals in Arabidopsis. Science 322, 1650–1655. 10.1126/science.1165594.

27. Williamson, R.E. (1991). Orientation of Cortical Microtubules in Interphase Plant Cells. Int Rev Cytol 129, 135–206.

28. Trinh, D.C., Alonso-Serra, J., Asaoka, M., Colin, L., Cortes, M., Malivert, A., Takatani, S., Zhao, F., Traas, J., Trehin, C., et al. (2021). How Mechanical Forces Shape Plant Organs. Preprint at Cell Press, 10.1016/j.cub.2020.12.001 https://doi.org/10.1016/j.cub.2020.12.001.

29. Saltini, M., and Deinum, E.E. (2025). Microtubule-based nucleation results in a large sensitivity to cell geometry of the plant cortical array. PLoS Comput Biol 21. 10.1371/journal.pcbi.1013282.

30. Kutschera, U., and Niklas, K.J. (2007). The epidermal-growth-control theory of stem elongation: An old and a new perspective. J Plant Physiol 164, 1395–1409. 10.1016/j.jplph.2007.08.002.

31. Savaldi-Goldstein, S., Peto, C., and Chory, J. (2007). The epidermis both drives and restricts plant shoot growth. Nature 446, 199–202. 10.1038/nature05618.

32. Coen, E., and Cosgrove, D.J. (2023). The mechanics of plant morphogenesis. Science (1979) 379. 10.1126/science.ade8055.

33. Landrein, B., and Hamant, O. (2013). How mechanical stress controls microtubule behavior and morphogenesis in plants: History, experiments and revisited theories. Plant Journal 75, 324–338. 10.1111/tpj.12188.

34. Muratov, A., and Baulin, V.A. (2015). Mechanism of dynamic reorientation of cortical microtubules due to mechanical stress. Biophys Chem 207, 82–89. 10.1016/j.bpc.2015.09.004.

35. Hamant, O., Heisler, M.G., Jönsson, H., Krupinski, P., Uyttewaal, M., Bokov, P., Corson, F., Sahlin, P., Boudaoud, A., Meyerowitz, E.M., et al. Developmental Patterning by Mechanical Signals in Arabidopsis.

36. Shaw, S.L., Kamyar, R., and Ehrhardt, D.W. (2003). Sustained Microtubule Treadmilling in Arabidopsis Cortical Arrays. Science (1979) 300, 1715–1718. 10.1126/science.1083529.

37. Nakamura, M., and Hashimoto, T. (2009). A mutation in the Arabidopsis gamma-tubulin-containing complex causes helical growth and abnormal microtubule branching. J Cell Sci 122, 2208–2217. 10.1242/jcs.044131.

38. Buschmann, H., Hauptmann, M., Niessing, D., Lloyd, C.W., and Schäffner, A.R. (2009). Helical growth of the Arabidopsis mutant tortifolia2 does not depend on cell division patterns but involves handed twisting of isolated cells. Plant Cell 21, 2090–2106. 10.1105/tpc.108.061242.

39. Chan, J., Sambade, A., Calder, G., and Lloyd, C. (2009). Arabidopsis cortical microtubules are initiated along, as well as branching from, existing microtubules. The Plant Cell Online 21, 2298–2306. 10.1105/tpc.109.069716.

40. Liu, T., Tian, J., Wang, G., Yu, Y., Wang, C., Ma, Y., Zhang, X., Xia, G., Liu, B., and Kong, Z. (2014). Augmin triggers microtubule-dependent microtubule nucleation in interphase plant cells. Current Biology 24, 2708–2713. 10.1016/j.cub.2014.09.053.

41. Nakamura, M., Ehrhardt, D.W., and Hashimoto, T. (2010). Microtubule and katanin-dependent dynamics of microtubule nucleation complexes in the acentrosomal Arabidopsis cortical array. Nat Cell Biol 12, 1064–1070. 10.1038/ncb2110.

42. Murata, T., Sonobe, S., Baskin, T.I., Hyodo, S., Hasezawa, S., Nagata, T., Horio, T., and Hasebe, M. (2005). Microtubule-dependent microtubule nucleation based on recruitment of gamma-tubulin in higher plants. Nat Cell Biol 7, 961–968. 10.1038/ncb1306.

43. Wasteneys, G. 0, and Williamson, R.E. (1989). Reassembly of microtubules in Nitella tasmanica: assembly of cortical microtubules in branching clusters and its relevance to steady-state microtubule assembly.

44. Thawani, A., Stone, H.A., Shaevitz, J.W., and Petry, S. (2019). Spatiotemporal organization of branched microtubule networks. Elife 8. 10.7554/eLife.43890.

45. Lindeboom, J.J., Nakamura, M., Hibbel, A., Shundyak, K., Gutierrez, R., Ketelaar, T., Emons, A.M.C., Mulder, B.M., Kirik, V., and Ehrhardt, D.W. (2013). A mechanism for reorientation of cortical microtubule arrays driven by microtubule severing. Science (1979) 342. 10.1126/science.1245533.

46. Chakrabortty, B., Willemsen, V., de Zeeuw, T., Liao, C.Y., Weijers, D., Mulder, B., and Scheres, B. (2018). A Plausible Microtubule-Based Mechanism for Cell Division Orientation in Plant Embryogenesis. Current Biology 28, 3031–3043.e2. 10.1016/j.cub.2018.07.025.

47. Kirchhelle, C., Chow, C.M., Foucart, C., Neto, H., Stierhof, Y.D., Kalde, M., Walton, C., Fricker, M., Smith, R.S., J??rusalem, A., et al. (2016). The Specification of Geometric Edges by a Plant Rab GTPase Is an Essential Cell-Patterning Principle During Organogenesis in Arabidopsis. Dev Cell 36, 386–400. 10.1016/j.devcel.2016.01.020.

48. Dong, J., MacAlister, C.A., and Bergmann, D.C. (2009). BASL Controls Asymmetric Cell Division in Arabidopsis. Cell 137, 1320–1330. 10.1016/j.cell.2009.04.018.

49. Mansfield, C., Newman, J.L., Olsson, T.S.G., Hartley, M., Chan, J., Mansfield, C., Newman, J.L., Olsson, T.S.G., Hartley, M., Chan, J., et al. (2018). Ectopic BASL Reveals Tissue Cell Polarity throughout Leaf Development in Arabidopsis thaliana Ectopic BASL Reveals Tissue Cell Polarity throughout Leaf Development in Arabidopsis thaliana. Current Biology 28, 2638–2646.e4. 10.1016/j.cub.2018.06.019.

50. Kuchen, E.E., Samantha, F., Barbier de Reuille, P., Kennaway, R., Bensmihen, S., Avondo, J., Calder, G.M., Southam, P., Robinson, S., Bangham, A., et al. (2012). Generation of Leaf Shape Through Early Patterns of Growth and Tissue Polarity. Science (1979) 335, 1092–1095. 10.1126/science.1214678.

51. Wallner, E.S., Mair, A., Handler, D., McWhite, C., Xu, S.L., Dolan, L., and Bergmann, D.C. (2024). Spatially resolved proteomics of the Arabidopsis stomatal lineage identifies polarity complexes for cell divisions and stomatal pores. Dev Cell 59, 1096–1109.e5. 10.1016/j.devcel.2024.03.001.

52. Yoshida, S., van der Schuren, A., van Dop, M., van Galen, L., Saiga, S., Adibi, M., Möller, B., ten Hove, C.A., Marhavy, P., Smith, R., et al. (2019). A SOSEKI-based coordinate system interprets global polarity cues in Arabidopsis. Preprint at Palgrave Macmillan Ltd., 10.1038/s41477-019-0363-6 https://doi.org/10.1038/s41477-019-0363-6.

53. Derbyshire, P., Findlay, K., McCann, M.C., and Roberts, K. (2007). Cell elongation in Arabidopsis hypocotyls involves dynamic changes in cell wall thickness. J Exp Bot 58, 2079– 2089. 10.1093/jxb/erm074.

54. Kutschera, U. (2008). The growing outer epidermal wall: Design and physiological role of a composite structure. Ann Bot 101, 615–621. 10.1093/aob/mcn015.

55. Verger, S., Long, Y., Boudaoud, A., and Hamant, O. (2018). A tension-adhesion feedback loop in plant epidermis. Elife 7. 10.7554/eLife.34460.

56. Tsitkov, S., Rodriguez, J.B., Bassir Kazeruni, N.M., Sweet, M., Nitta, T., and Hess, H. (2022). The rate of microtubule breaking increases exponentially with curvature. Sci Rep 12, 20899. 10.1038/s41598-022-24912-0.

57. Kent, I.A., and Lele, T.P. (2017). Microtubule-based force generation. Preprint at Wiley-Blackwell, 10.1002/wnan.1428 https://doi.org/10.1002/wnan.1428.

58. Aher, A., Rai, D., Schaedel, L., Gaillard, J., John, K., Liu, Q., Altelaar, M., Blanchoin, L., Thery, M., and Akhmanova, A. (2020). CLASP Mediates Microtubule Repair by Restricting Lattice Damage and Regulating Tubulin Incorporation. Current Biology 30, 2175–2183.e6. 10.1016/j.cub.2020.03.070.

59. Zhang, Y., Iakovidis, M., and Costa, S. (2016). Control of patterns of symmetric cell division in the epidermal and cortical tissues of the Arabidopsis root. Development. 10.1242/dev.129502.

60. Panteris, E., Adamakis, I.D.S., Daras, G., and Rigas, S. (2015). Cortical microtubule patterning in roots of Arabidopsis thaliana primary cell wall mutants reveals the bidirectional interplay with cell expansion. Plant Signal Behav 10. 10.1080/15592324.2015.1028701.

61. Lindeboom, J.J., Nakamura, M., Hibbel, A., Shundyak, K., Gutierrez, R., Ketelaar, T., Emons, A.M.C., Mulder, B.M., Kirik, V., and Ehrhardt, D.W. (2013). A mechanism for reorientation of cortical microtubule arrays driven by microtubule severing. Science (1979) 342, 1245533. 10.1126/science.1245533.

62. Sambade, A., Pratap, A., Buschmann, H., Morris, R.J., and Lloyd, C. (2012). The influence of light on microtubule dynamics and alignment in the Arabidopsis hypocotyl. Plant Cell 24, 192–201. 10.1105/tpc.111.093849.

63. Vineyard, L., Elliott, A., Dhingra, S., Lucas, J.R., and Shaw, S.L. (2013). Progressive transverse microtubule array organization in hormone-induced Arabidopsis hypocotyl cells. Plant Cell 25, 662–676. 10.1105/tpc.112.107326.

64. Atkinson, S., Kirik, A., and Kirik, V. (2014). Microtubule array reorientation in response to hormones does not involve changes in microtubule nucleation modes at the periclinal cell surface. J Exp Bot 65, 5867–5875. 10.1093/jxb/eru325.

65. Kennaway, R., Coen, E., Green, A., and Bangham, A. (2011). Generation of diverse biological forms through combinatorial interactions between tissue polarity and growth. PLoS Comput Biol 7. 10.1371/journal.pcbi.1002071.

66. Kennaway, R., and Coen, E. (2019). Volumetric finite-element modelling of biological growth. Open Biol 9. 10.1098/rsob.190057.

67. Mitchison, T., and Kirschner, M. (1984). Dynamic instability of microtubule growth. Nature 312, 237–242. 10.1038/312237a0.

68. Chan, J., Calder, G., Fox, S., and Lloyd, C. (2007). Cortical microtubule arrays undergo rotary movements in Arabidopsis hypocotyl epidermal cells. Nat Cell Biol 9, 171–175. 10.1038/ncb1533.

69. Calder, G., Hindle, C., Chan, J., and Shaw, P. (2015). An optical imaging chamber for viewing living plant cells and tissues at high resolution for extended periods. Plant Methods 11, 22. 10.1186/s13007-015-0065-7.

70. Ueda, K., Matsuyama, T., and Hashimoto, T. (1999). Visualization of microtubules in living cells of transgenicArabidopsis thaliana. Protoplasma 206, 201–206. 10.1007/BF01279267.

71. Chan, J., Calder, G., Fox, S., Lloyd, C. (2005). Localization of the Microtubule End Binding Protein EB1 Reveals Alternative Pathways of Spindle Development in Arabidopsis Suspension Cells. the Plant Cell Online 17, 1737–1748. 10.1105/tpc.105.032615.

72. Abe, T., and Hashimoto, T. (2005). Altered microtubule dynamics by expression of modified *α* -tubulin protein causes right-handed helical growth in transgenic Arabidopsis plants. The Plant Journal 43, 191–204. 10.1111/j.1365-313X.2005.02442.x.

73. Chan, J., and Coen, E. (2020). Interaction between Autonomous and Microtubule Guidance Systems Controls Cellulose Synthase Trajectories. Current Biology 30, 941–947.e2. 10.1016/j.cub.2019.12.066.

74. ImageJ Https://imagej.nih.gov/ij/. Preprint.

75. Thévenaz, P., Ruttimann, U.E., and Unser, M. (1998). A pyramid approach to subpixel registration based on intensity. IEEE Transactions on Image Processing 7, 27–41. 10.1109/83.650848.

76. Fonck, E., Feigl, G.G., Fasel, J., Sage, D., Unser, M., Rüfenacht, D.A., and Stergiopulos, N. (2009). Effect of aging on elastin functionality in human cerebral arteries. Stroke 40, 2552– 2556. 10.1161/STROKEAHA.108.528091.

