## Supplemental Data for "Tissue-wide cues are sensed at the cellular level to coordinate microtubule orientations in plants"

Supplementary Material

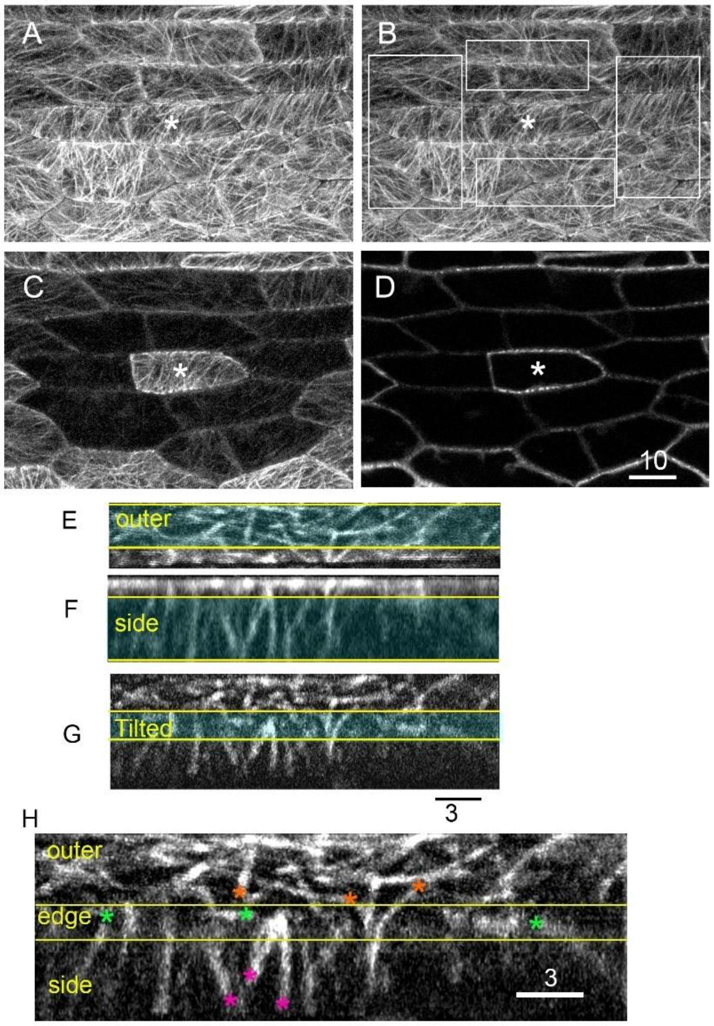

**Figure S1. Imaging single fluorescent cells.**

(**A-D**) Generating a single fluorescent cell by photo-bleaching its neighbours. (**A**) Outer epidermal faces of leaf cells expressing TUA6-GFP before bleaching. (**B**) Bleaching was performed by targeting the laser onto the cortical regions marked by the boxes. To prevent bleaching of the cell of interest (marked by the asterisk) the boxes avoided its edges. (**C, D**) Outer epidermal faces and mid planes through cells after 10 min exposure to the laser. Microtubules appear as spots in transverse sections as they are aligned vertically on side faces. When specifically bleached, fluorescence is strong in the cell of interest but not between interfaces of bleached cells. Double-headed arrow in A indicates orientation of the leaf proximodistal axis. Bar = 10 µm

(**E-H**) Unfolded views of cells from 3D time-lapse data .Three views projected from the z-stacks near an edge of a single fluorescent cell: planar view of the outer face (**E**), planar view of the side face (**F**), and tilted view with both faces (**G**). (**H**) Combined unfolded view based on cyan boxes with yellow edges of E-G. The edge domain captured from the tilted section and is shown between yellow lines. Asterisks indicate examples of microtubule plus ends on the outer face (orange), in the edge domain (green) and the side face (magenta). Bar = 3 µm.

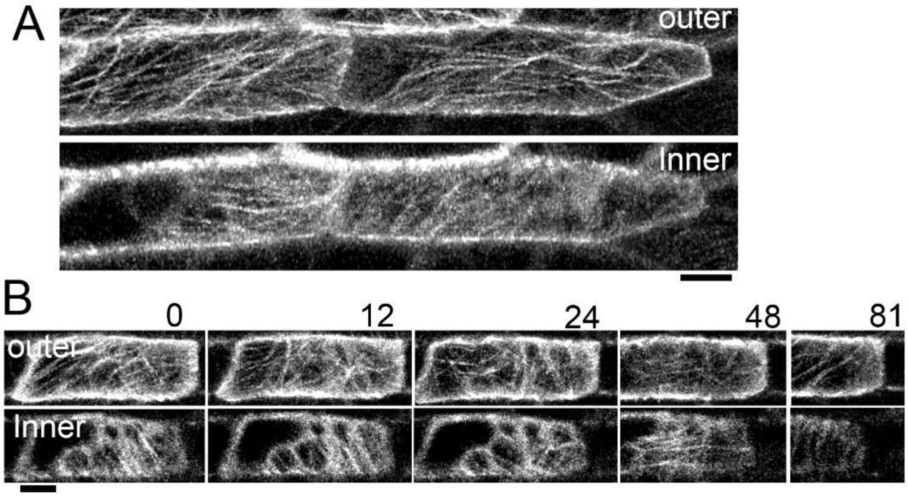

**Figure S2. Persistent reorientation on inner and outer faces of adjacent or single fluorescent cells**

(**A**) Projections of confocal z-stack show microtubule alignments on inner faces (bottom panel) can differ from the outer faces (top panel). These single fluorescent cells were prepared by photobleaching the fluorescent signal of microtubules in only the cells neighbouring one side. Bar = 5 µm

(**B**) Projections of movie frames showing microtubule alignments on inner faces are dynamic. The alignment on the inner face changes from oblique (0 min) to longitudinal (48 min) and then becomes transverse (81 min). The cell moves out of the imaging field due to growth of the leaf. Bar = 5 µm.

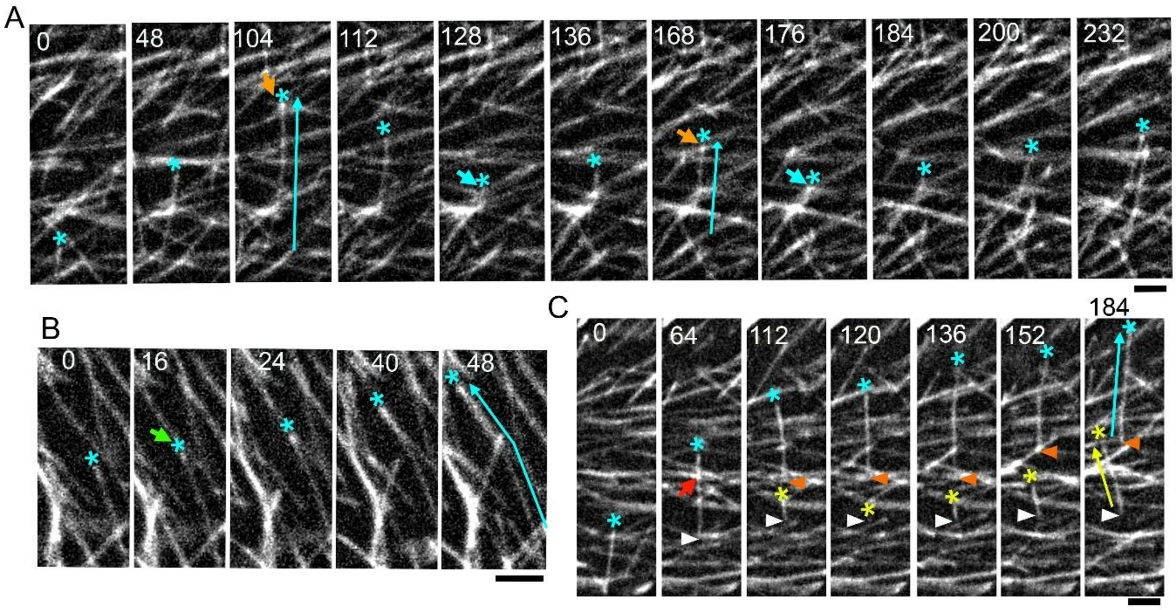

**Figure S3. Examples of catastrophe, rescue, zipper and severing on outer cell face**

(**A**) Time-lapse series showing collision-induced catastrophe and rescue at crossover junctions. A growing plus end (cyan asterisk) crosses several microtubules before undergoing catastrophe (orange arrow) upon collision with an obstructing microtubule at a steep angle (104 sec). The shrinking end is rescued (cyan arrow) at a crossover junction (128 sec). The microtubule undergoes another cycle of collision-induced catastrophe (168 sec) and rescue at a crossover junction (176 sec). Cyan arrows indicate the trajectories of the microtubule upon catastrophe. Bar = 1.5 µm.

(**B**) Time-lapse series showing zippering. A growing plus end (cyan asterisk) encounters an obstructing microtubule at a shallow angle (green arrow, 16 sec). Its trajectory is redirected along the path of the obstructing microtubule. The cyan arrow indicates the microtubule's final trajectory, which is kinked due to zippering. Bar = 2 µm.

(**C**) Time-lapse series showing sever-rescue. A growing plus end (cyan asterisk) crosses several obstructing microtubules. The red arrow (64 sec) indicates the crossover junction where severing will take place. Severing results in development of a gap between the newly exposed minus (orange arrowhead) and plus ends (yellow asterisk, 112-120 sec), as the latter undergoes rapid depolymerisation. The new plus end is recued close to its minus end (white arrowhead) and generates a new trajectory. Cyan and yellow arrows indicate the trajectories of the leading and lagging pieces of the severed microtubule. Bar = 2 µm.

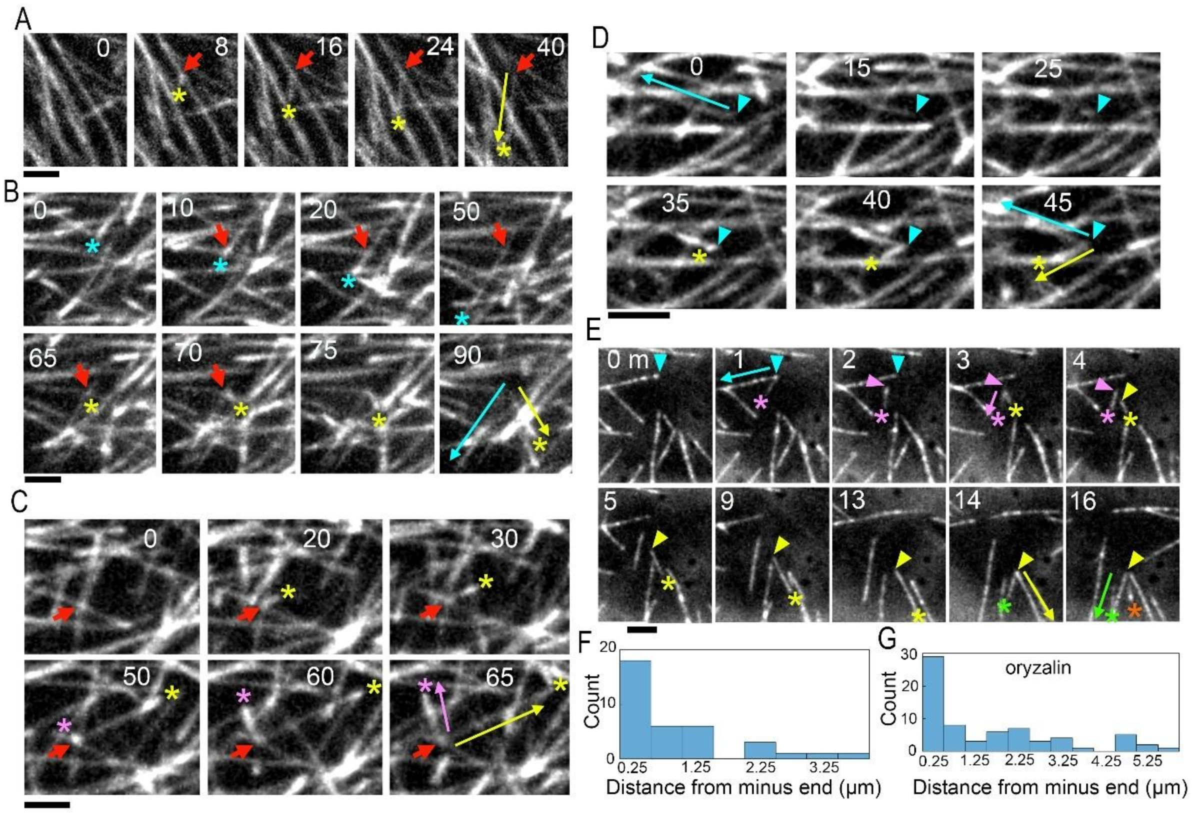

**Figure S4. Examples of microtubule branching on outer cell face**

(**A**) Movie frames showing initiation of a free branch from the side of an extant microtubule. Red arrow indicates the site where branch appears. The yellow asterisk marks the growing plus end of the young branch. Yellow arrow indicates the final trajectory of the branch. Time in seconds. Bar = 1.5 µm.

(**B**) Movie frames showing initiation of a crossover branch. Red arrow indicates the crossover junction from where the branch appears. The crossover junction was made by the growing plus end marked by the cyan asterisk (10). The yellow asterisk marks the plus end of the branch emerging from the intersection (65). Cyan and yellow arrows indicate the final trajectories of the crossing microtubule and branch, respectively. Bar = 1.5 µm.

(**C**) Movie frames showing branching from an intersection. Red arow indicates an intersection from where branching occurs. Yellow asterisk indicates the initiation of the first branch from the intersection. Magenta asterisk indicates a second branch from the intersection. Yellow and magenta arrows indicate the final trajectories of the branches. Bar = 1.5 µm.

(**D**) Movie frames showing branching from close to minus end of parent microtubule. Minus end of parent microtubules (formed after severing). The yellow asterisk marks the plus end of a branch that initiates from close to the minus end (indicated with cyan arrowhead). Yellow and cyan arrows in final timepoint (45) indicate the final trajectories of the branch and its parent. Bar =2.0 µm.

(**E**) Movie frames showing branching close to the minus ends of microtubules in cells treated with oryzalin. Successive branches initiate from the minus ends of daughter microtubules. Arrows indicate the trajectories of parent microtubules in frames when branches emerge from close to minus ends of a parent microtubules (1 min, 3 min, 14 min, 16 min). The minus ends of mother microtubules and the plus ends of branching microtubules are marked by arrowheads and asterisks, respectively. The orange asterisk marks the emergence of another branch from the axil of the green trajectory (16 min). Bar =2.0 µm.

(**F-G**) Frequency histograms of branching distances from the parent minus end in untreated cells (**F**) n=36 branches, 3 cells, 3 plants and in cells treated with oryzalin (**G**), n=69 branches, 5 cells, 2 plants. Midpoint of alternate 0.5 µm-wide bins are numbered.

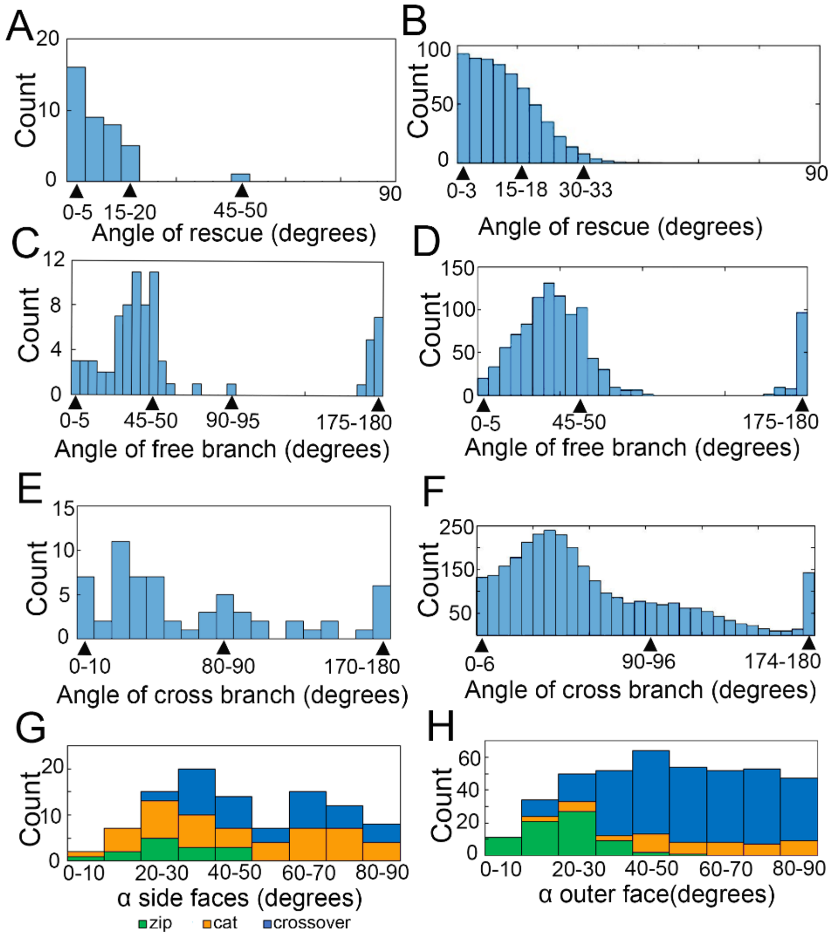

### Figure S5. Measured angles compared to outputs of ground state model or to other faces

(**A-B**) Frequency histograms of measured rescue angles on outer face, n= 3 cells, 39 rescues (**A**) compared with those from the output of ground-state model (**B**). Arrow heads indicate bin values.

(**C-D**) Frequency histograms of measured free branching angles on outer face relative to the parent microtubule, n= 3 cells, 77 free branches (**C**) compared with those from the output ground-state model (**D**).

(**E-F**) Frequency histograms of measured crossover-induced branch angles on outer face relative to the crossing microtubule, n= 3 cells, 62 crossover branches (**E**), compared with those from the output of ground-state model (**F**).

(**G-H**) Frequency histograms of measured angles (α) of redirection (green), catastrophe (orange) and crossover (blue) on side faces 100 encounters, 6 cells (**G**), compared with the outer face, 417 encounters, 9 cells (**H**).

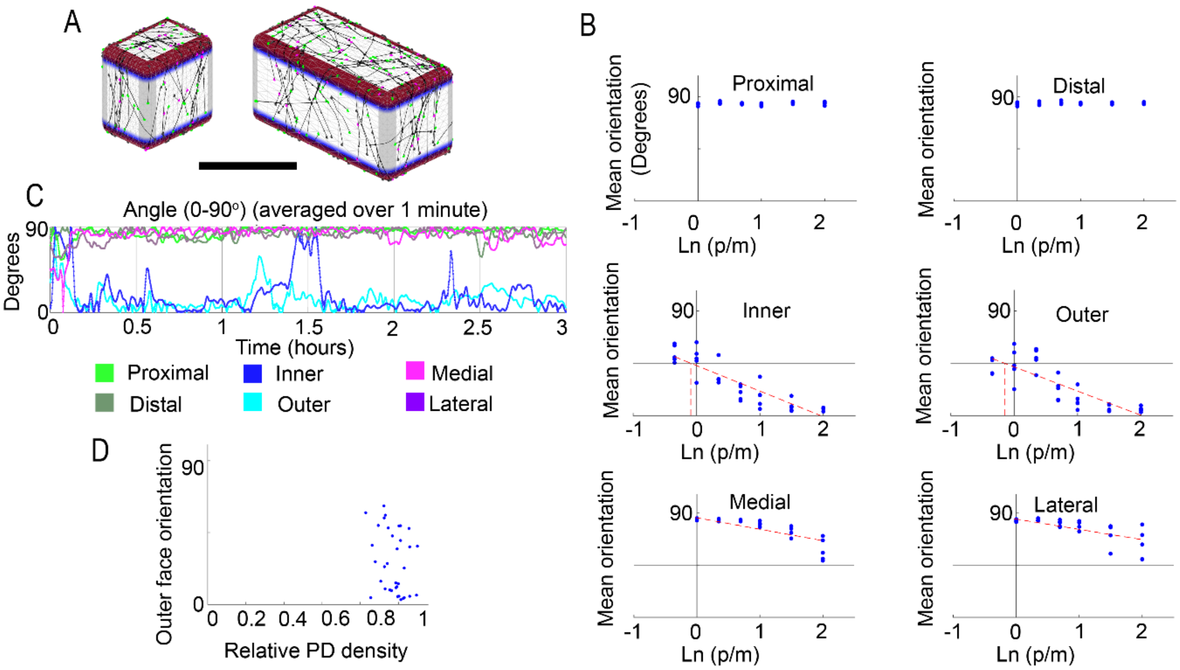

### Figure S6. Output of model that combines edge-induced catastrophe, edge generation and orthogonal edge-filter

(**A**) Microtubule pattern at end of runs in cells with p/m = 0.7 or 2 Ln aspect. Grey = Edge catastrophe, blue = edge-orthogonal filter, red= edge generation. Note red and grey overlap to give dull red. Bar = 10 μm.

(**B**) Mean microtubule orientations on faces of cells plotted against ln (p/m). Alignments display long axis geometric bias on outer and inner faces. All side faces display short axis geometric bias (Proximal, Distal, Medial and Lateral). 3 runs per cell aspect. (line fit equations: Inner: y=-22x+43, R2=0.78; Outer: y=-21x+42, R2=0.69; Medial: y=-9.9x+86, R2=0.62; Lateral: y=-8.9x+85, R2=0.52 ).

(**C**). Mean microtubule angle plotted against time. Outer face shows persistent reorientation. Plot from cell with p/m = 2.

(**D**) Mean outer face orientation for different runs and values of p/m, plotted against density of microtubules on PD faces divided by the side face mean.

#### Table S1. Parameters measured in outer epidermal leaf cell face

| **Parameter (Outer face)** | **Experimental measurement** |
|---|---|
| Plus-end density (growing plus ends per $\mu m^{-2}$) | 0.32 (SD = 0.07; n= 3 cells, 3 plants) |
| Microtubule length (μm) | 5 (SD=0.58, n=3 cells, 3 plants, 87 microtubules) |
| Plus-end growth rate ($\mu m\ s^{-1}$) | 0.09 (SD = 0.01; n=3 cells, 3 plants, 90 plus ends) |
| Plus-end shrink rate ($\mu m\ s^{-1}$) | 0.22 (SD = 0.03, n=3 cells, 3 plants, 33 plus ends) |
| Plus-end spontaneous rescue rate ($s^{-1}$of shrink) | 0.04 (n= 3 cells, 3 plants,19 rescues, 31 shrinking ends over a period of 501 sec of shrink). |
| Plus-end rescue probability at crossover junctions | 0.13 (n=1 cell, 1 plant, 12 rescues, 91 encounters) |
| Rescue probability for new plus end after severing | 0.03 (n= 3 cells, 3 plants, 55 severing events) |
| Rescue probability for new minus end after severing | 1 (n= 3 cells, 3 plants, 55 severing events) |
| Probability of crossover-induced severing | 0.16 (n= 3 cells, 3 plants, 56 severing events, 368 crossovers) |
| Crossover severing delay (s) | 63 (SD =13; n= 3 cells, 3 plants, 54 severing events) |
| Probability the crossing microtubule is severed | 0.7 (54 severing events, 3 cells, 3 plants). |
| Free branching rate ($s^{-1}$per μm of microtubule) | 0.0002 (SD=4.1E-05, n= 3 cells, 3 plants) |
| Free branching rate ($s^{-1}$per microtubule) | 0.001 (SD = 0.0001, n= 3 cells, 3 plants) |
| Branch sever delay (s) | 71 (SD = 17; 11 branches, 3 cells, 3 plants) |
| Probability of crossover branch | 0.08 (n= 3 cells, 3 plants, 27 crossover branches, 361 crossovers) |
| Crossover branch delay (s) | 60 (SD = 30; n= 3 cells, 3 plants, 27 branches) |
| Probability of branch arising from the crossing microtubule | 0.4 (1 cell, 1 plant) |
| Probability of edge induced catastrophe on OI edges | 0.7 (n=3 cells, 3 plants, 17 catastrophes, 25 microtubules) |
| Probability of edge induced catastrophe on non-OI edges | 1 (n= 3 cells, 3 plants, 43 catastrophes, 43 microtubules) |
| Plus-end emergence rate from ML edges ($s^{-1}$per μm of edge) | 0.36 (SD= 0.07, 4 cells, 2 plants; over a period of 792 sec) |
| Plus-end emergence rates from PD edges ($s^{-1}$per μm of edge) | 0.14 (SD= 0.05, 4 cells, 2 plants; over a period of 792 sec) |
| Rescue angles | See Fig. S5A (n=3 cells, 3 plants, 39 rescues) |
| Free branching angles | See Fig. S5C (n= 3 cells, 3 plants, 77 branches) |
| Crossover branch angles | See Fig. S5E (n=3 cells, 3 plants, 62 branches) |
| Minus-end shrink rate ($\mu m\ s^{-1}$) | 0.008 (n= 3 cells, 3 plants, 34 minus ends) |
| MT-MT interaction probabilities | See Fig. S5H (n=9 cells, 9 plants, 417 encounters) |

#### Table S2. Parameters measured in outer epidermal leaf cell face and one of the side faces in single-fluorescent cells

| Parameter | Outer face (strip) | ML faces | PD faces |
|---|---|---|---|
| Plus-end density (growing plus end per μm$^{-2}$) | 0.14 (SD = 0.06; n= 4 cells, 4 plants) | NA | 0.05 (SD = 0.02; n=4 cells, 4 plants) |
| Plus-end growth rate (μm s$^{-1}$) | 0.08 (SD = 0.02; n=3 cells, 3 plants, 28 plus ends) | NA | 0.09 (SD = 0.09; n=3 cells, 3 plants, 22 plus ends) |
| Plus-end shrink rate (μm s$^{-1}$) | 0.18 (SD=0.02, n=3 cells, 3 plants, 30 plus ends) | NA | 0.23 (SD=0.03, n=3 cells, 3 plants, 21 plus ends) |
| Plus-end rescue rate (s$^{-1}$ of shrink) | 0.03 (SD = 0.008, n=3 cells, 3 plants, 29 shrinking ends) | NA | 0.02 (SD=0.004, n=3 cells, 3 plants, 21shrinking ends) |
| Plus-end spontaneous catastrophe rate (s$^{-1}$of growth) | | | 0.003 (SD=0.001; n=3 cells, 3 plants, 32 plus ends) |
| Free branching rate (s$^{-1}$per growing plus end) | 0.001 (SD=0.001; n=4 cells, 4 plants, 8 branches) | NA | 0 (n=4 cells, 4 plants) |
| Minus-end shrink rate (μm s$^{-1}$) | 0.01 (SD = 0.001; n= 2 cells, 2 plants, 6 minus ends) | NA | 0.01 (SD = 0.005; n= 2 cells, 2 plants, 6 minus ends) |
| Plus-end density (growing plus end per μm$^{-2}$) | 0.51 (SD = 0.08; n= 3 cells, 3 plants) | 0.24 (SD = 0.09; n= 3 cells, 3 plants) | NA |
| Plus-end growth rate (μm s$^{-1}$) | 0.09 (SD = 0.003; n=2 cells, 2 plants, 29 plus ends) | 0.1 (SD = 0.01; n=2 cells, 2 plants, 28 plus ends) | NA |
| Plus-end shrink rate (μm s$^{-1}$) | 0.24 (SD=0.003, n=2 cells, 2 plants, 27 plus ends) | 0.26 (SD=0.03, n=2 cells, 2 plants, 25 plus ends) | NA |
| Plus-end rescue rate (s$^{-1}$of shrink) | 0.03 (SD = 0.009, n=2 cells, 2 plants, 27 shrinking ends) | 0.03 (SD=0.01, n=2 cells, 2 plants, 26 shrinking ends) | NA |
| Plus-end spontaneous catastrophe rate (s$^{-1}$of growth) | | 0.002 (SD=9.5705E-05; n=2 cells, 2 plants, 43 plus ends) | NA |
| Free branching rate (s$^{-1}$per microtubule) | 0.0015 (SD=0.0004; n=3 cells, 3 plants, 51 branches) | 0.0001 (SD=0.0001; n=3 cells, 3 plants, 2 branches) | NA |
| Minus-end shrink rate (μm s$^{-1}$) | NA | 0.03 (SD = 0.03; n= 1 cell, 1 plant, 2 minus ends) | NA |

| | | | |
|---|---|---|---|
| Interaction probabilities (PD and ML faces) | | Fig. S5G (n=6 cells, 6 plants, 100 encounters) | Fig. S5G (n=6 cells, 6 plants, 100 encounters) |

Parameters were measured on the outer and one of the side faces (either PD or ML) of single fluorescent cells. Imaging was limited to 2 faces due to the constraint imposed by imaging side faces within 2-3 seconds so that microtubules could be tracked.

#### Table S3. Parameters used in models

| Parameter name in model | Value | Description |
|---|---|---|
| **Cell properties** | | |
| Lr = 'YXlogratio' | 1 | Cell shape: log of the aspect ratio (0 for a cube) |
| **Microtubule properties** | | |
| Dm = 'tubulethickness' | 0.025 | Microtubule diameter (μm) |
| rg+ = 'plus_growthrate' | 0.09 | Plus-end growth rate (μm $s^{-1}$) |
| rs- = 'minus_shrinkrate' | 0.008 | Minus-end shrink rate (μm $s^{-1}$) |
| rd = 'plus_shrinkrate' | 0.2 | Plus-end shrink rate (μm $s^{-1}$) |
| psc = 'prob_plus_catastrophe' | 0.0015 | Plus-end spontaneous catastrophe probability ($s^{-1}$of growth) |

| | | |
|---|---|---|
| pr = 'prob_plus_rescue' | 0.02 | Plus-end rescue probability unless crossover ($s^{-1}$of shrink) |
| pcr = 'prob_crossover_rescue' | 0.13 | Plus-end rescue probability if a shrinking end crosses another microtubule |
| **Rescue angle profile** | | |
| αr = 'rescue_angle_mean' | 0.1745 | Mean rescue angle in radians (10 degrees) |
| σr = 'rescue_angle_spread' | 0.1745 | Mean spread of rescue angle in radians (10 degrees) |
| **Limits on microtubule density** | | |
| magh = 'max_growing_mt_per_area' | 1.0 | Maximum density of growing plus ends per unit area |
| miρ = min_mt_density | 0.01 | Minimum density of microtubules, measured as total microtubule area/total cell area |
| ρrs = density_min_rescue_sharpness | 2 | How rapidly the rescue rate increases with drop in density (Arbitrary units) |
| **Free Nucleation/Branching** | | |
| ic = 'initialcreationperarea' | 0.3 | Number of initial microtubules ($\mu m^{-2}$) |
| pfb = 'prob_tail_branch_time' | 0.001 | Free branching probability ($s^{-1}$per minus end) |
| **Branch Angle profiles** | | |
| pbf = 'prob_branch_forwards' | 0.89 | Probability that a new branch is generally in the forwards direction of its parent |
| αf = 'branch_forwards_mean' | 0.6981 | Mean of branch angle in radians, for forward branch angles (40 degrees) |
| σf = 'branch_forwards_spread' | 0.2967 | Std dev in radians of absolute branch angle for forward branch angles (17 degrees) |

| | | |
|---|---|---|
| pbp = 'prob_branch_antiparallel' | 0.8182 | For branching generally in the backwards direction relative to the parent, probability exactly antiparallel to it |
| αb = 'branch_backwards_mean' | 0.1745 | Mean of branch angle in radians, for backward branch angles (10 degrees) |
| σb = 'branch_backwards_spread' | 0.0873 | Std dev in radians of absolute branch angle, for backward branches (5 degrees) |
| **Local microtubule-microtubule encounter rules** | | |
| αvs, αs 'collision_angles' | [0.3491 0.6981] | Collision angle upper bounds (20° and 40° in radians) |
| pz = 'probs_zip' | [0.7, 0.4, 0] | Probability of zippering for angles <20°, 20 °-40°, 40°-90° |
| pc = 'probs_cat' | [0.1, 0.1, 0.2] | Probability of catastrophe for angles <20°, 20°-40°, 40°-90°. |
| ps = 'prob_crossover_cut' | 0.16 | Probability of crossover severing. |
| psc = 'prob_crossover_cut_collider' | 0.7 | Probability the crossing microtubule is severed. |
| pcb = 'prob_crossover_branch' | 0.08 | Probability of crossover branch |
| pob = 'prob_crossover_branch_collided' | 0.3 | Probability of branch arising from the obstructing microtubule |
| **Delays** | | |
| ds = 'crossover_sever_delay' | 60 | Crossover severing delay (s) |
| db = 'crossover_branch_delay' | 60 | Crossover branch delay (s) |
| **Edge-induced catastrophe** | | |
| pce = 'cat_per_edge' | 0.7 or 0 | Probability that a plus end entering an edge at right angles catastrophises before exiting the region (set to 0 for ground state model) |

| | | |
|---|---|---|
| **Stress alignment** | | |
| fs = 'field_alignment' | -0.06 or 0 | Strength of a vertically aligned field on side faces, set to 0 except for stress models (Arbitrary units) |
| foi = 'field_alignment2' | -0.006 or 0 | Strength of a PD aligned field on outer and inner faces, set to 0 except for stress models (Arbitrary units) |
| **Edge generation** | | |
| pe = 'edgecreate_perareasecond' | 0.025 or 0 | Probability of generating new microtubules in the edge regions ($s^{-1}$per $\mu m^{-2}$). Set to 0 except for differential face models |
| spd = 'edgecreaterate_pd_scale' | 0.5 or n/a | Scaling of probability of edge creation on horizontal PD edges |
| **Edge-orthogonal filter** | | |
| αi = 'barrier_incidence' | 50° | Angle of incidence above which plus ends undergo catastrophe when they arrive at the edge-orthogonal filter |

### Measurements of local microtubule behaviours and model input parameters

GFtbox is a Matlab package for modelling the three-dimensional growth of biological tissues, which can be downloaded from https://github.com/JIC-Enrico-Coen/GrowthToolbox. Tissues are modelled by decomposing the volume into polyhedra and applying finite element methods to numerically solve the differential equations of growth and elasticity. Method and software are detailed in [65,66] .

Here we have added the capability of modelling microtubules growing along the inner surface of a cell wall. The cell is modelled as a cuboid with rounded edges and spherical corners. The shape is approximated in GFtbox by a mesh of triangles, each given a constant thickness (making them triangular prisms). The GFtbox project for the model in this paper can be accessed at https://github.com/JIC-Enrico-Coen/Microtubules_in_growing_leaves_2025. In situations where we need to know the curvature of the surface at some point, we use the known curvature of the continuous shape rather than attempting to estimate it from the finite elements, which are themselves only an approximation to that shape. This avoids compounding approximations. Examples of situations where this arises are extending a microtubule across a rounded edge following a geodesic path.

### Cell properties

Cell dimensions in the imaged epidermal region were measured using line tool of ImageJ. Cell lengths along PD (indicated by 'p' in Figure 1B) and ML ('m' in Figure 1B) axes were measured at the start and end of movies to calculate average lengths. The average value for p and m were 47 µm and 19 µm, respectively (n = 82 cells, 2 leaves). For the model, cells were 10 µm x 10 µm x *p* µm, where *p* was the length along the proximodistal axis (Figure 1B) and varied from 5µm to 74µm to allow cells with different p/m values to be modelled. Curvature at edges and corners had a radius of 1 µm.

### Microtubule growth and shrink rates

Parameters for the outer face were based on manually tracking microtubules in movies of 3 cells (total of 90 microtubules). Growth and shrink rates were determined by dividing length grown or shrunk by time spent in growth or shrink, respectively. For plus ends, a mean growth rate was estimated to be 0.09 µm per sec (total of 90 microtubules), and a mean shrink rate during catastrophe of 0.2 µm per sec (total of 33 shrinking plus ends). Minus ends predominantly underwent shrink. A mean minus end shrink rate was estimated to be 0.008 µm per sec (total of 34 minus ends). Minus end growth was ignored. These values were used to set model parameters for plus end growth rate, $r_{g+}$, plus end shrink rate during catastrophe, $r_d$, and minus end shrink rate, $r_{s-}$, respectively. Microtubules were modelled assuming a diameter $D_m$ = 0.025 µm, based on electron microscopy images[3].

### Rescue probabilities and angles

Conversion of a shrinking plus end back to growth is known as rescue[67]. Measurements from shrinking ends were divided into 2 groups depending on their origin: (1) from plus end collisions with other microtubules or induced in the edge region and (2) from severing. Rescue rates were determined by dividing the number of rescue events by time spent in shrink. A mean rescue rate of 0.04 events per plus end per sec of shrink was obtained for plus ends shrinking after collisions with other microtubules (3 cells, total of 19 rescues, 31shrinking ends, 501 secs of shrink). A similar rate was obtained for plus ends formed after severing (0.03, 17 rescues, 59 shrinking ends, 547 secs of shrink).

Sometimes, the location of rescue coincided with a crossover junction (27 out of 44 rescues, 3 cells; see Figure S3A). This behaviour was quantified in a movie with high resolution - all rescue events were examined. The probability of rescue at a junction was determined by dividing the number of rescues at a junction by total number of junctions encountered by shrinking end, which gave a value of 0.13 (12 rescues at junction, 91 encounters). A spontaneous rescue rate of 0.02 per plus end per sec of shrink was estimated for microtubules undergoing rescue in junction-free spaces (9 rescues, 41 shrinking ends, 455 secs of shrink). Based on these findings, probability of a plus end undergoing spontaneous rescue, $p_r$, was set to 0.02 per plus end per sec, and probability that a shrinking plus end undergoes rescue when crossing a microtubule, $p_{cr}$, was set to 0.13 per crossover.

The distribution of rescue angles, measured relative to original trajectory, was measured for from tracking individual microtubules in 3 cells (Figure S5A). Based on these results, angle distribution for both spontaneous and crossover-induced rescue was modelled assuming a rescue angle mean, $\alpha_r$ = 10°, with a standard deviation $\sigma_r$ = 10°. The resulting distribution of angles from a simulation is shown in Figure S5B.

### Microtubule plus-end density

The plus end density (per μm$^2$) was measured by counting the total number of plus-end comets in a snapshot of a cell face and dividing that number by the cell face area. This gave a value of 0.32 for the outer face.

### Limits on microtubule density

To prevent runaway increase in microtubule density, there was an upper limit on the number of growing microtubule heads per μm$^2$ of $ma_{gh}$ = 1. This upper limit was implemented by reducing the probability of creating a new growing head as the number of existing ones approaches the limit. Whenever the simulation requests the creation of a growing head (which may happen either because of a microtubule branching or being severed at a crossover, or the spontaneous creation of a new microtubule) the probability that it is created is 1 – (number of growing heads)/($ma_{gh}$ × cell area), or zero if that is negative. If creation fails, the branch or new microtubule is not started, or the rear half of the severed microtubule immediately catastrophises.

To prevent irreversible stochastic loss of microtubules, there is a lower limit on microtubule density $mi_\rho$ = 1% of the cell face area. This was implemented by increasing the probability per unit time that a catastrophising microtubule undergoes rescue as the lower limit is approached. The probability is scaled by a factor which is close to 1 when the ratio of current density to $mi_\rho$ is well above 1, and increases as the density ratio falls towards 1. If we define $mi_{\rho f}$, the minimum density fraction, as (current density)/$mi_\rho$ (or 1 if that is larger), then the probability per unit time of a rescue is multiplied by $1 + \rho_{rs} /(mi_\rho - 1)$. $\rho_{rs}$ is a parameter that determines how fast the probability scaling increases as $mi_{\rho f}$ falls towards 1. The smaller this value, the sharper the increase. When $mi_{\rho f} < 1$, rescue probability is 1 and thus rescue following catastrophe is instantaneous.

### Branching rates

Branching is reported to occur along microtubules and at intersections formed by microtubules crossing each other[39,40,42]. We therefore scored branching along microtubules (free branching: Figure S4A) and at intersections from movies (crossover branching: Fig S4B, C). We identified these events through direct observation. Care was taken to ensure the new plus end was not associated with rescue of either the mother microtubule, or the crossing and obstructing microtubules. The formation of a gap at intersections was used to distinguish branching from severing at intersections. Some crossover branches may have arisen by sever-rescue if the new plus end was rescued immediately after severing.

Branches that arose from microtubules without intersection were classed as free branches. To see if free branching occurred uniformly along microtubules, we measured the distances they initiated from the minus end using measure tool of ImageJ. Microtubules in bundles were not included as the locations of minus ends of parent microtubules were not clear. Free branching was biased towards the minus end of the parent microtubule (Figure S4D, E).

Branching angle was measured relative to the plus end of the mother microtubule or the crossing microtubule in the cases of crossover branches using the Angle tool of ImageJ. Free branching rate from a microtubule was estimated using the following calculation: The free branching rate per μm of microtubule per sec is equal to the number of new free branches/number of plus ends in movie frame x average length of microtubules/time observed in secs.

Average length of microtubules was estimated to be 5 μm from measurements of single microtubules, where both ends could be seen (3 cells, 87 microtubules). They were measured upon interaction with other microtubules (i.e. catastrophe or zipper) or with edges. These measures gave an average free branching rate of 0.00021 per μm of microtubule per sec (3 cells; 0.00024. 0.00016, 0.00023).

Free branching rate was also estimated per microtubule per sec by calculating the number of new free branches/number of plus ends/duration of movie in seconds. This calculation gave a mean branch rate of 0.001 branches per microtubule per sec (3 cells, 0.0012, 0.0009, 0.001).

Crossover branching rate was estimated in the same way as free branching rate, giving a cross-over branching rate of 0.0008 per μm of microtubule per sec (3 cells; 0.001, 0.0007, 0.0006). Thus crossover-induced branching rate is similar to free branching rate on the outer face.

Models were seeded at the beginning of each simulation with $i_c$ = 0.3 per $\mu m^2$ randomly oriented and positioned microtubule plus ends. The probability of free branching from the minus end, $p_{fb}$, was set to 0.001 per sec in accordance with the observed rate.

### Free branch angle profiles

The observed distribution of angles for free branching, relative to the direction of the parent microtubule is shown in Figure S5C. This distribution was captured in the model by assuming the probability that a branch was generally in the forward direction relative to the parent, $p_{fp}$, was 0.89 (and thus a generally backwards probability of 0.11). For generally forwards branches, the mean branch angle, $\alpha_f$ , was set to 40° and the standard deviation, $\sigma_f$ to 17°. For the generally backwards direction, the probability that the branch was exactly antiparallel to the parent was $p_{bp}$ = 0.8182, the mean branch angle, $\alpha_b$, was set to 10° and the standard deviation, $\sigma_b$, to 5°. Branches were produced to the left or right of the parent with equal probability. The resulting distribution for a typical simulation is shown in Figure S5D.

### Microtubule-microtubule encounters

When the growing plus end of a microtubule encounters an obstructing microtubule, different outcomes may arise with different probabilities depending on the angle of encounter, α. The frequencies of zippering, catastrophe and crossover (Figure S5G and S5H) were measured by manually tracking the outcomes of encounters between plus ends and obstructing microtubules. Angles were measured using the Angle tool of ImageJ.

Probabilities of crossover, zippering and catastrophe for three angle intervals ($<\alpha_{vs}$, $\alpha_{vs}$ - $\alpha_s$° and $> \alpha_s$), where $\alpha_{vs} = 20°$ and $\alpha_s = 40°$, were estimated based on observed fractions of each event on the outer face (Figure 2A). These values were used to set the parameters in the decision tree of Figure 2B. Probability of zippering $p_z = 0.7$ for $\alpha < \alpha_{vs}$, $p_z = 0.4$ for $\alpha_{vs} < \alpha < \alpha_s$, and $p_z = 0$ for $\alpha > \alpha_s$. Similarly, we assigned a probability of catastrophe, $p_c = 0.1$ for $\alpha < \alpha_{vs}$, $p_c = 0.1$ for $\alpha_{vs} < \alpha < \alpha_s$, and $p_c = 0.2$ for $\alpha > \alpha_s$. Probability of crossover for each angle interval was $1 – p_z - p_c$.

We also measured probabilities of severing or branching following a crossover by dividing the number of severing or branching events by the number of crossovers. Crossover junctions less than 60 sec old in a movie were not included since it was found that it took about this time for severing or branching to be observable after crossover (means of 63 and 60 sec, respectively; 3 cells). Based on these findings we assigned a probability of severing, $p_s = 0.16$ (mean of 3 cells: 0.14, 0.20, 0.14) and a probability of crossover-induced branching of $p_{cb} = 0.08$ (mean of 3 cells: 0.13, 0.07, 0.04).

If severing occurred, we found a probability of cutting the crossing microtubule of 0.7 (mean of 3 cells: 0.6, 0.8, 0.7). We therefore set the probability that the crossing microtubule was severed, $p_{sc}$, to 0.7. After severing, the new plus end underwent catastrophe, while the new minus end was stable. The mean rescue rate of the new plus end was determined to be 0.03 per sec (3 cells, total of 17 rescues, 59 shrinking plus ends, 547 sec shrink), similar to the rate for crossover-induced catastrophes.

To capture the distribution of crossover branching angles (Fig S5E), we assumed that branching angle was determined as for free branching but was oriented relative to the obstructing microtubule with a probability $p_{ob} = 0.3$. The mean delay between crossover and severing or branching was 63 and 60 sec, respectively (3 cells), so we incorporated a delay of 60 secs for each from the time of crossover in the model (parameters $d_s$ and $d_b$). Based on these assumptions the distribution of crossover branch angles from the simulation output was similar to that observed (compare Fig S5E with S5F).

To illustrate interaction rules, we simulated them for one or two microtubules (Movie 3).

### Edge-induced catastrophe

The edge region was defined to be where the cell surface is curved at edges and corners. For models including edge-induced catastrophe, the probability of a plus end undergoing catastrophe was elevated while plus end resided within the edge region.

Microtubules entering the edge regions at lower angles therefore had a higher probability of catastrophe before exiting the edge region because of their greater dwell time. $p_{ce}$ = 0.7, was the probability that a tubule entering an edge at right angles catastrophises before exiting the far side. This probability was converted to a catastrophe rate per unit time while the plus end is within an edge region, which was added to the base rate of catastrophe per unit time everywhere in the edges and corners.

### Stress-sensing

The stress field was represented using the orientations of a vector field on each face. (Unlike a vector, a stress does not have a sign associated with. Strictly speaking, stress is represented with a nematic tensor field rather than a vector field, but a vector field was sufficient in this case for modelling purposes). The effect of the stress field on a growing tubule head was assumed to induce a curvature of the growing region tending to reduce the angle between the growth direction and the stress field. This tended to bring the tubule into either parallel or antiparallel alignment with the stress field. The induced curvature was zero when the plus end growth direction was parallel, antiparallel, or perpendicular to the vector field. A suitable function with this property was sin(2θ), where θ is the angle from the growth direction to the axis of the stress field. This is multiplied by a constant. There are two such constants, $f_s$ = field alignment for an orthoplanar stress field on the side faces, and $f_{oi}$ = field alignment for a mediolateral stress field on the outer and inner faces. The magnitudes of these parameters were chosen to the be the minimum needed to capture the observed microtubule alignment patterns. Values of these parameters were 0 in all models except the stress-cue model.

### Edge generation

Edge-generated microtubules had a uniform distribution of angle probabilities. Probability of generation (nucleation) from horizontal (OI) edge regions per unit of area per second was set by $p_e$ = 0.025. To generate ML bias, the probability of edge generation was scaled by $s_{pd}$ =0.5 on PD edges.

### Edge-orthogonal filter

The edge-orthogonal filter was specified to be at the border of the O and I faces, where they abut side faces. Microtubules arriving at the filter underwent catastrophe if their angle of incidence was greater than $a_i$, set to 50°.

#### Model output measures

### Mean angle

The mean angle of tubules on a face at a given time is calculated by finding the best-fit ellipse to the set of directions of all tubule segments, weighted by their lengths. The unsigned angle between the main axis of this ellipse and a reference direction for each face is the mean angle. It is in the range 0 to π/2 since the sense of the reference axis and the sense of the tubule growth direction are both ignored. For the aspect-angle plots, these means are then averaged over the whole run (omitting a warm-up period of 30 minutes at the start).

#### Side face densities

The density of tubules in a face is the total length of all tubule segments within that face, multiplied by the constant tubule thickness, divided by the face area. Relative PD densities were calculated from the mean densities on the PD faces divided by the average of the densities on the ML and PD faces.
